# Dynamic cortical behavior of plant protoplasts reveals unexpected similarities between plant and animal cells

**DOI:** 10.1101/2024.11.13.623369

**Authors:** Johanna E. M. Dickmann, Marjolaine Martin, Claire Lionnet, Zoe Nemec-Venza, Olivier Hamant

## Abstract

In contrast to animal cells with their contractile cortical cytoskeleton, plant cells are encased in walls and are usually thought to behave like stiff, pressurized vessels. When digesting the cell wall, plant protoplasts form spherical shapes, further consolidating this model. Here we challenge this apparent opposition between animal and plant cells. We show that plant protoplasts can form protrusions. More precisely, using plasma membrane markers, we reveal that 27 ± 1% of the protoplasts show dot-like protrusions and 16 ± 7% of protoplasts show long (typically 1-20 μm, up to 200 μm) protrusions that we name “filopods”. We demonstrate that this behavior is independent of the plant species, the membrane marker or the osmolyte. Protrusions can host large and long structures, such as microtubule bundles, which in turn can impact the mechanical behavior of the protrusions. We find that filopods can form *de novo* when increasing osmolarity, that they most likely move passively, and that they can attach to artificial surfaces. Altogether, these observations show that forming and retaining protrusions is not exclusive to animal cells. This calls for revisiting the dynamics and probing ability of the plant cell cortex in different osmotic and mechanical environments.

## Main

From a biophysical standpoint, plant cells can be caricatured as passive “bags of membrane”, pressed against a rigid cell wall by turgor pressure. Indeed, turgor pressure would burst the plasma membrane in the absence of the load-bearing cell wall. In contrast, animal cells do not have a cell wall and are mainly shaped by a cytoskeletal cell cortex that can actively deform the membrane, e.g. forming protrusions^1^. Yet, plant cells can also form protrusions. Indeed, when plants are exposed to a hyperosmotic environment, turgor pressure is released, water leaves the cell, and the protoplast (i.e. the cell without its wall) retracts from the cell wall in a process called plasmolysis ^2^. During this process, the plasma membrane stays connected to the cell wall via the Hechtian structure ^3,4,5,6^ (Fig. 1a), comprised of membranous tubes called Hechtian strands, a network-like structure close to the cell wall called the Hechtian reticulum, and discrete points connecting the plasma membrane to the cell wall, called Hechtian attachment sites. These protrusions appear as a consequence of pre-attachment sites on the wall and do not imply an autonomous ability of plant protoplasts to generate protrusions. This is what we explore here.

**Fig. 1.**
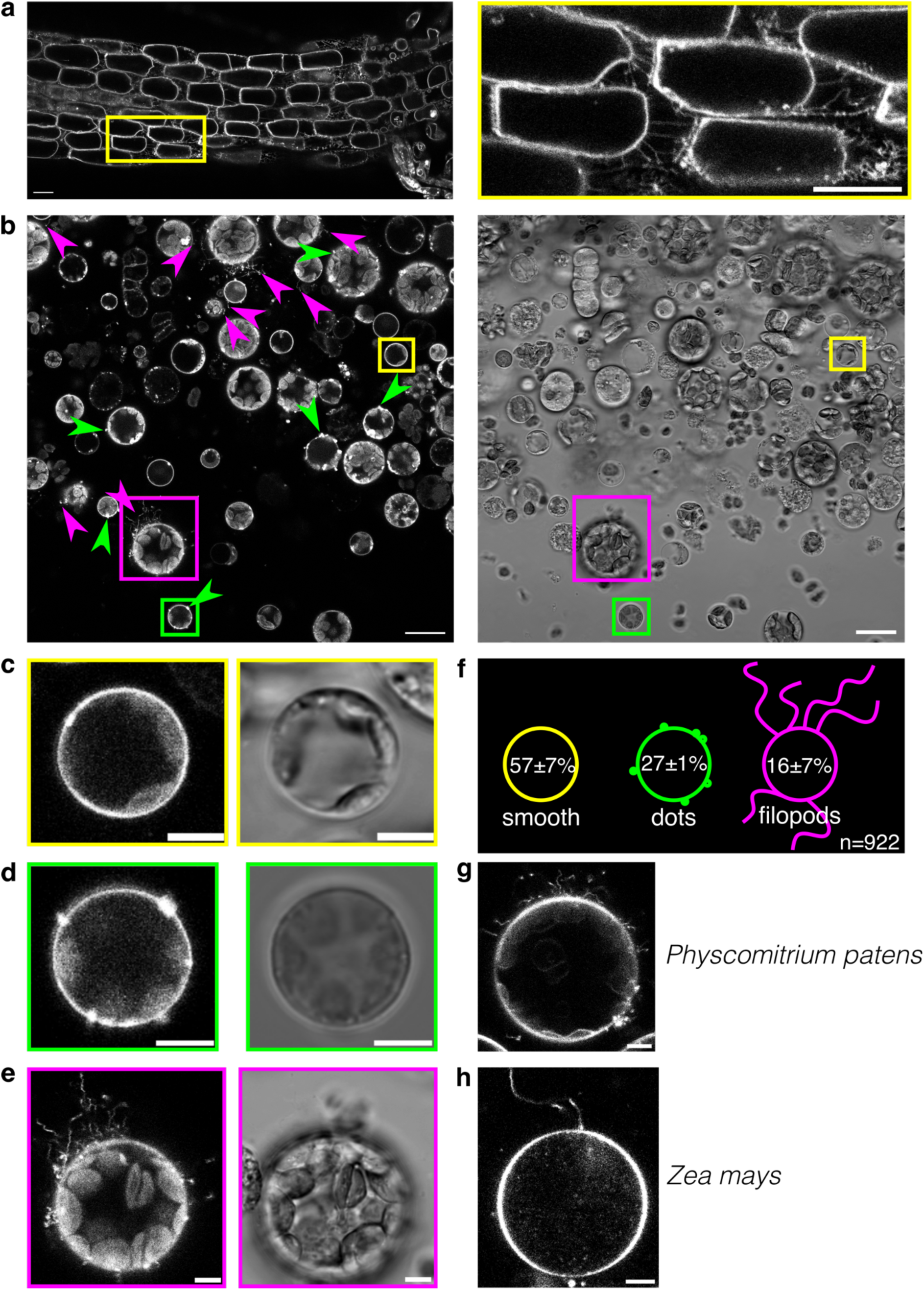
Plant cells exhibit protrusions upon release of turgor pressure and digestion of the cell wall. **a**, Plasma membrane marker (*pUBQ10::LTi6B-TdTomato*) observed in the epidermal cell layer of the Arabidopsis hypocotyl, plasmolyzed in a hyperosmotic buffer containing 600 mM D-mannitol (∼720 mOsmol/kg) for 50 min reveals Hechtian structures, i.e. membrane-encased protrusions connecting the protoplast to the cell wall. Left: Close up boxed in yellow. Scale bars: 20 µm. **b-e**, Arabidopsis protoplasts labelled by plasma membrane marker (*pUBQ10::LTi6B-TdTomato*) in NOA73 containers (ca. 4x4 mm, height: ca. 2 mm). Scale bars: 20 µm (**b**), 5 µm (**c-e**). **b**, Large scale field of view of protoplasts after digestion. **c-e**, Close-ups marked with respective colors in **b**. **c**, Smooth protoplast. **d**, Protoplast with short protrusions called “dots”; additional protoplasts with dots are marked with green arrow heads in **b**. **e**, Protoplasts with long protrusions called “filopods”; additional protoplasts with filopods are marked with magenta arrow heads in **b**. **f**, Relative numbers of occurrence of respective protoplast types based on N = 3 independent experiments with a total of n = 922 alive protoplasts. **g-h**, Protoplasts stained with membrane dye FM4-64, scale bars 5 µm. **g**, *Physcomitrium patens* (bryophyte), representative image of N = 3 independent experiments. **h**, *Zea mays* (monocotyledon), representative image of N = 3 independent experiments. All fluorescent images show single confocal planes.

### Protoplasts robustly form protrusions

When releasing the cell wall by digestion in a hyperosmotic buffer solution (Extended Data Fig. 1a), smooth protoplasts are observed (Fig. 1b,c). Taking a closer look at Arabidopsis protoplasts from whole seedlings using a plasma membrane marker line, we discovered that, to our surprise, many protoplasts exhibited membrane-encased protrusions (Fig. 1b,d,e). These took either the form of short, round protrusions that we named “dots” (27 ± 1% of all alive protoplasts, Fig. 1d,f, Table S1) or of long, thin protrusions that we termed “filopods” (16 ± 7% of all alive protoplasts, Fig. 1e,f, Table S1), as their shape recalls the shape of filopodia observed in animal cells. Typically, dots had a diameter between 0.5 and 3 µm and filopods had lengths between 1 and 20 µm. The longest filopods we could observe were up to 210 µm long (Extended Data Fig. 2).

We observed both kinds of protrusions in protoplast suspensions contained in custom made polymer (NOA73) containers (Fig. 1b-e), NOA73 microwells (see Fig. 2c, Fig. 3, Fig. 4, Fig. 5a-d) as well as between a glass slide and cover slip (Extended Data Fig. 3). Furthermore, protrusions were also observed when using sorbitol instead of mannitol as an osmolyte (Extended Data Fig. 1b). Last, beyond *Arabidopsis thaliana* (Fig. 1b-f), protrusions were readily observed in protoplasts obtained from the bryophyte *Physcomitrium patens* (Fig. 1g) as well as the monocotyledon *Zea mays* (Fig. 1h), when stained with the lipophilic dye FM4-64. Incidentally, this also shows that the formation of protrusions is independent from the fluorescent marker.

**Fig. 2.**
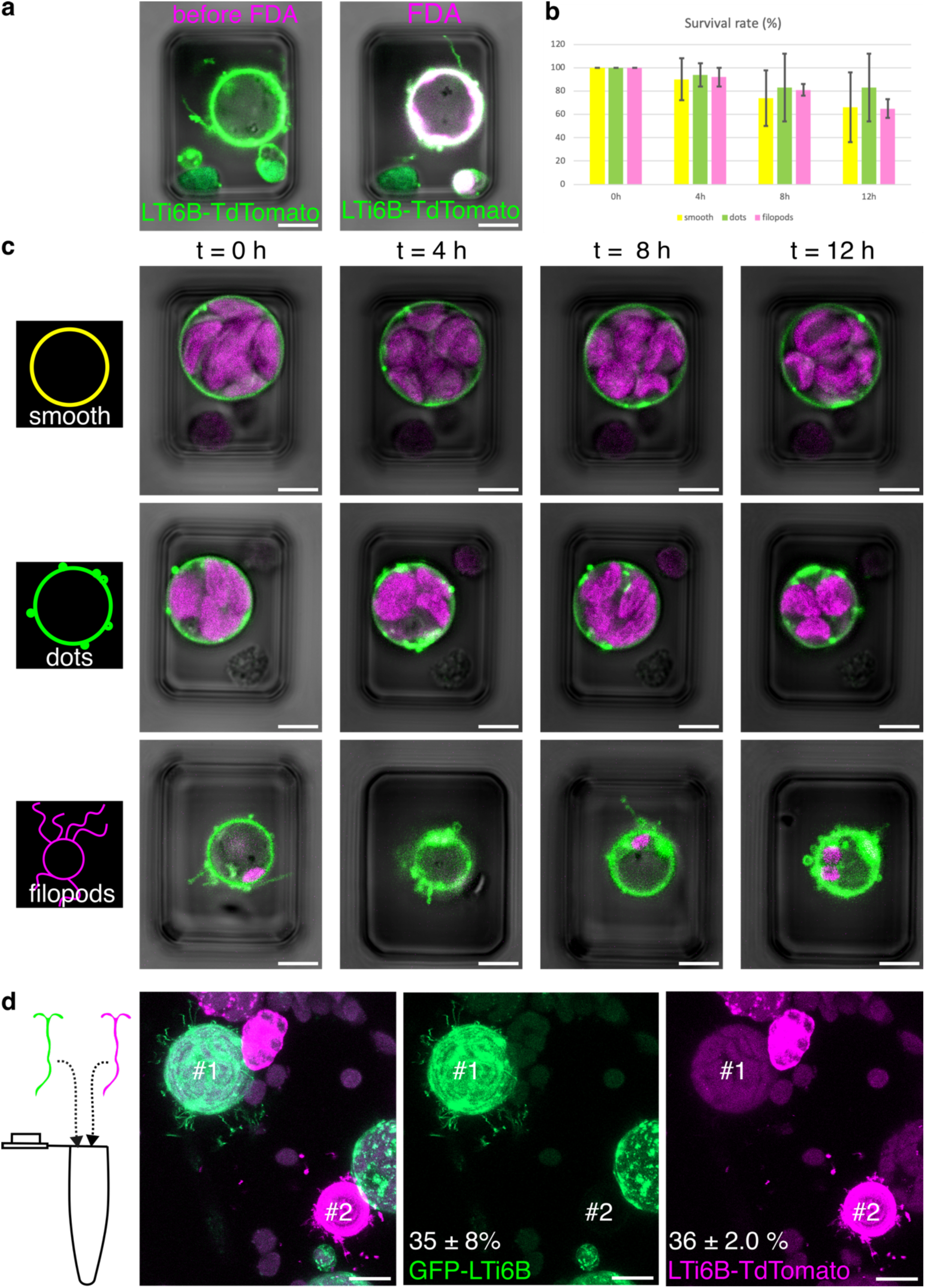
Protoplasts with protrusions are viable and the protrusions are continuous with the protoplast’s plasma membrane. **a**, Membrane marker (*pUBQ10::LTi6B-TdTomato*), pseudo-colored in green, shows protoplast with filopods in custom-made NOA73 microwells (15x20 µm, 21 µm high), overlaid with fluorescein signal, pseudo-colored in magenta, before (left) and after (right) addition of the vital stain fluorescein diacetate (FDA), and with transmitted light image. Fluorescent images: single confocal plane. **b**, Survival of protoplasts of different categories. The average survival rate of each category ± standard deviation is shown for each time point. Yellow: smooth, green: protoplasts with dots, magenta: protoplasts with filopods. **c**, Overnight time course imaging of smooth protoplasts, protoplasts with dots, and protoplasts with filopods demonstrates long-term protoplast viability. Time after onset of the timelapse is marked on top. The total number of protoplasts observed for the respective categories across N = 3 independent experiments are n = 43 (smooth), n = 15 (dots), n = 39 (filopods). The fluorescent images show single confocal planes. Overlay of membrane marker (*pUBQ10::LTi6B-TdTomato*), pseudo-colored in green, chloroplast autofluorescence, pseudo-colored in magenta, and transmitted light image. **d**, Protrusions are continuous with the plasma membrane. Left: Schematic shows joint digestion of plants from category *#*1 (*p35S::GFP-LTi6B)* and *#*2 (*pUBQ10::LTi6B-TdTomato)*. Center: Maximum intensity projection capturing the fluorescence of both *p35S::GFP-LTi6B* (labelled *#*1, pseudo-colored in green) and *pUBQ10::LTi6B-TdTomato* (labelled *#*2, pseudo-colored in magenta). Right: Individual channels. The numbers show the average percentages ± standard deviation of protoplasts with protrusions (sum of filopods and dots) of category #1 and #2, respectively (n = 574 alive protoplasts across N = 3 independent experiments). We did not find a single protoplast of category *#*1 with magenta protrusions nor a protoplast of category *#*2 with green protrusions. Scale bars: 5 µm (**a**, **c**), 10 µm (**d**).

**Fig. 3.**
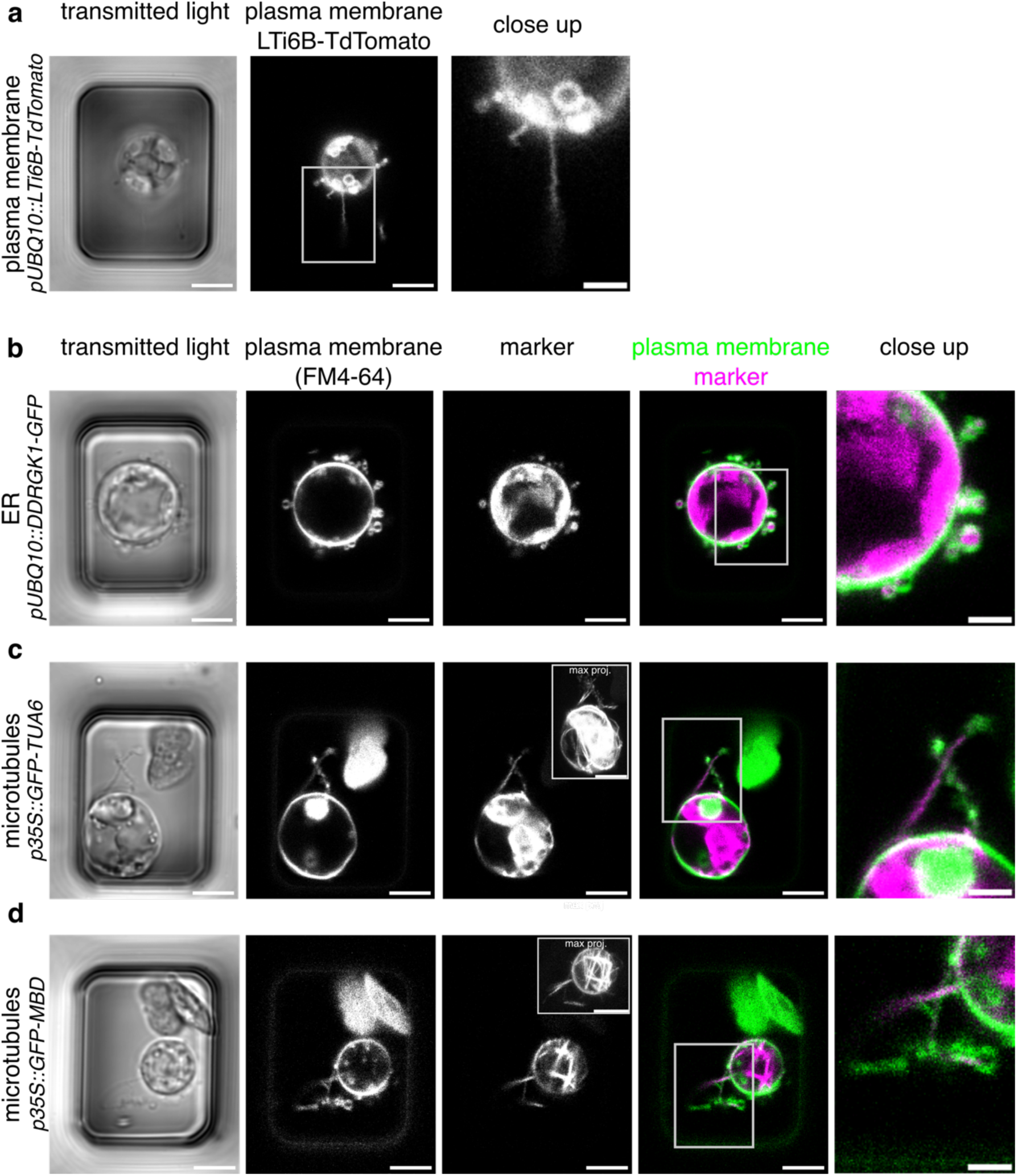
Protrusions can contain ER and microtubules. **a**, *pUBQ10::LTi6B-TdTomato* expressing protoplast (plasma membrane marker). **b***, pUBQ10::DDRGK1-GFP* expressing protoplast (ER membrane marker), plasma membrane is stained with FM4-64. **c**, *p35S::GFP-TUA6* expressing protoplast (tubulin marker), plasma membrane is stained with FM4-64. **d**, *p35S::GFP-MBD* expressing protoplast (microtubule bundle marker), plasma membrane is stained with FM4-64. Fluorescent images show single confocal planes; inset in marker images in **c**,**d** shows maximum intensity projection of whole z-stack. Close ups are marked by rectangles in precedent image. Scale bars: 5 µm, Close ups: 2 µm.

**Fig. 4.**
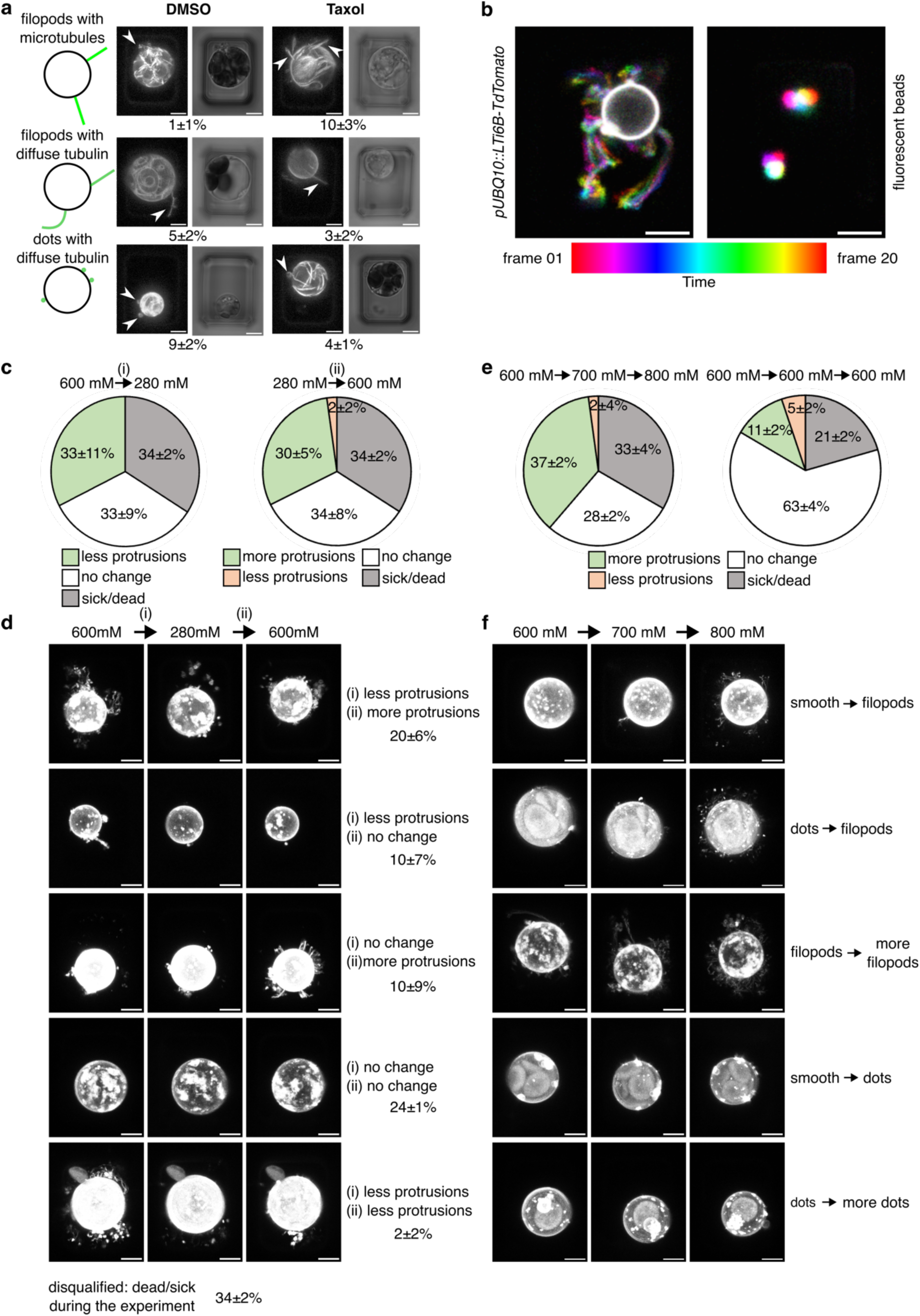
Dynamics and regulation of the protrusions. **a**, Microtubule stabilization by taxol induces more microtubule-containing, straight filopods. Sketches show the three possible configurations of protrusions detectable with a tubulin marker (*p35S::GFP-TUA6*). Top: straight filopods with strong GFP-TUA6 signal (indicating microtubule bundles). Middle: straight or bent filopods with diffuse GFP-TUA6 signal (indicating diffuse tubulin or individual microtubules). Bottom: dots with diffuse GFP-TUA6 signal. Images show examples for each category in control condition (DMSO) and after microtubule stabilization with taxol. Maximum intensity projection of the GFP-TUA6 signal (arrows indicate the respective protrusions, gamma = 0.5, see Extended Data Fig. 8b for images with gamma = 1), transmitted light image. Percentages state average fraction of protoplasts with the respective protrusions ± standard deviation over n = 298 (taxol) or n = 300 (DMSO) protoplasts across N = 3 independent experiments. **b**, The movement of the protrusions is qualitatively similar to the movement of fluorescent beads and hence may be passive. Timelapse of a single confocal plane of a protoplast with a plasma membrane marker (*pUBQ10::LTi6B-TdTomato*) (left) compared to fluorescent polystyrene beads (right), both in NOA73 microwells. 20 consecutive time frames (frame time 103ms) were color-coded and superposed to visualize movement, see also Video S2. Images representative for N = 3 independent experiments. **c**, **d**, The protrusions are partially taken up in low osmolarity and re-formed in higher osmolarity. Protoplasts with plasma membrane marker (*pUBQ10::LTi6B-TdTomato*) were subjected to changes in protoplasts medium: (i) decrease in osmolarity from 600 mM D-mannitol (720 mOsmol/kg) to 280 mM D-mannitol (340 mOsmol/kg), (ii) increase in osmolarity from 280 mM D-mannitol back to 600 mM D-mannitol. Quantification of the behavior of n = 97 protoplasts across N = 3 independent experiments: average ± standard deviation for **c**, each change of medium separately or **d**, both changes combined and example images. Maximum intensity projections of plasma membrane marker. **e**,**f**, New protrusions are formed with increasing osmolyte concentration. Protoplasts with a plasma membrane marker (*pUBQ10::LTi6B-TdTomato*) were subjected to an increase in osmolarity in the protoplast medium: 600 mM D-mannitol ∼ 720 mOsmol/kg to 700 mM D-Mannitol ∼ 850 mOsmol/kg to 800 mM D-mannitol ∼ 980 mOsmol/kg. **e**, Quantification of the protoplast behavior (n = 75 protoplasts across N = 3 independent experiments, left), compared to control (n = 97 protoplasts across N = 3 independent experiments, right, where medium of the same osmolarity was added). Green: more protrusions, i.e. smooth protoplasts developing dots or filopods, protoplasts with dots developing more/bigger dots or filopods, protoplasts with filopods increasing the number and/or length of their filopods. Red: less protrusions, i.e. protoplasts with dots showing less dots, protoplasts with filopods showing less filopods or only dots. White: no change, i.e. smooth protoplasts staying smooth, protoplasts with dots keeping a similar number and size of dots, protoplasts with filopods keeping a similar number and size of filopods. **f**, Examples of the different options of the green category as specified on the right. Maximum intensity projections of plasma membrane marker (*pUBQ10::LTi6B-TdTomato*). Scale bars: 5 µm.

**Fig. 5.**
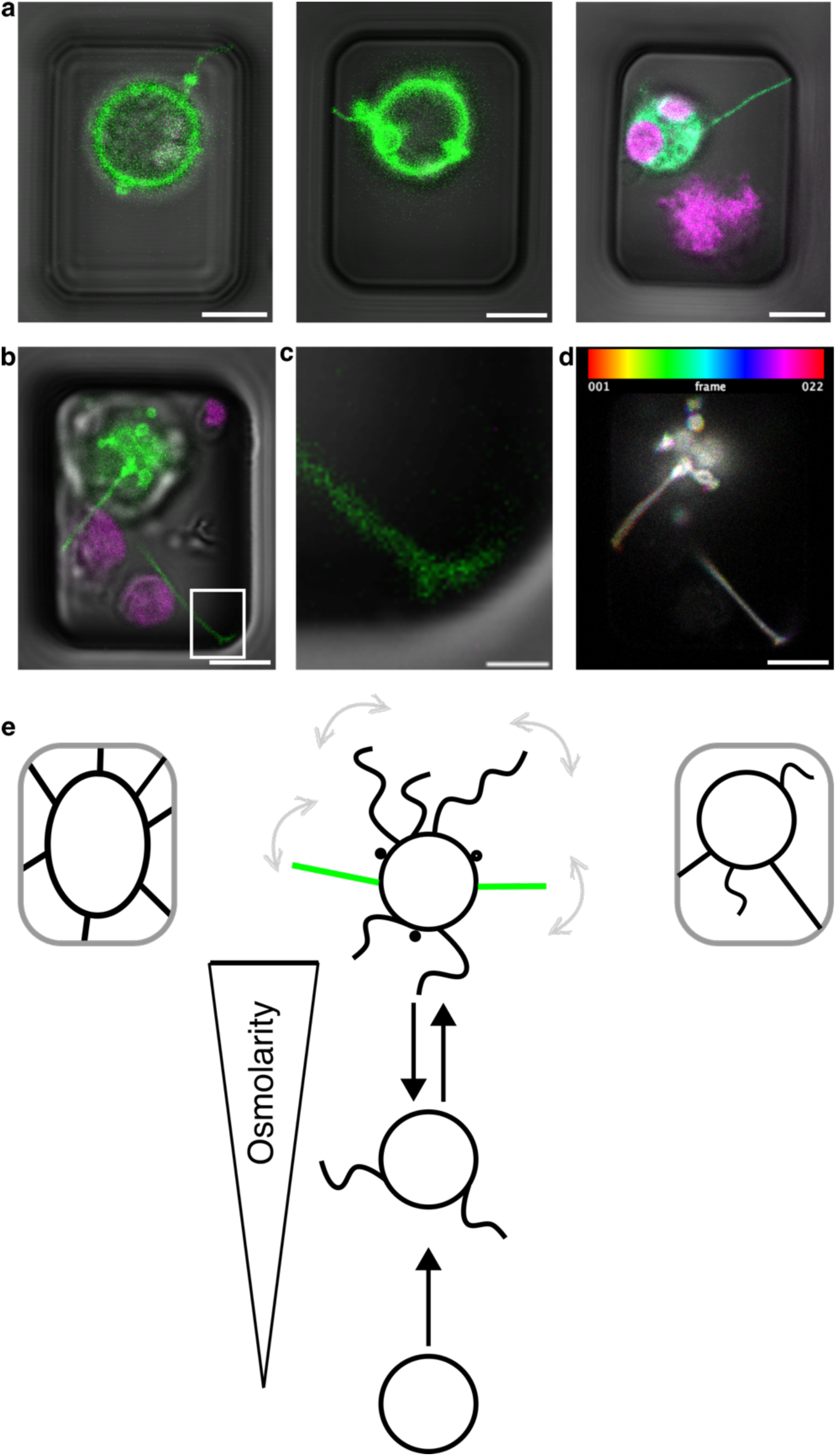
**a,** Filopods are occasionally observed to attach to the surface of the microwells. Example images from two different experiments: left, middle: plasma membrane marker (*pUBQ10::LTi6B-TdTomato*), pseudo-colored in green; right: tubulin marker (*p35S::GFP-TUA6*), pseudo-colored in green, each overlayed with transmitted light (note the microwell) and chloroplast autofluorescence (pseudo-colored in magenta). See also movies S3-5. b-d, Example of a filopod contacting the microwell and widening the contact area. b, plasma membrane marker (*pUBQ10::LTi6B-TdTomato*) pseudo colored in green, overlayed with chloroplast autofluorescence (pseudo-colored in magenta) and transmitted light. c, magnification of area boxed in b: widening of filopod tip at contact to the wall of the microwell. d, Timelapse movie of the protoplast shown in b, time is color-coded as specified by the time scale bar on top, frame time 630ms. White shows overlay between timepoints and thus absence of movement. Scale bars: 5 µm (a, b, d), 1 µm (c). e, Plant cells show protrusions in hyperosmotic environment when they plasmolyze while the plasma membrane (black) stays connected to the cell wall (grey) via the Hechtian structure (left). Protoplasts also show protrusions that can take the shape of dots, filopods, or straight, microtubule-containing filopods (green) in the absence of cell wall (middle). These protrusions are regulated by osmolarity (bottom): The length and density of filopods is reversibly increased with increasing osmolarity. Smooth protoplasts can form *de novo* protrusions with increasing osmolarity. Protoplasts can contact the walls of microwells (grey) they are contained in with their filopods, suggesting that filopods could form *de novo* Hechtian attachment sites, engaging in a “search and capture” mechanism (right).

### Protrusions originate from the plasma membrane of living protoplasts

Surprised by this observation, we wondered whether the protrusions were a sign of cell death. In order to test for protoplast viability, we used vital stains. To image the same protoplasts before and after staining, protoplasts were contained in microwells^7^. We used two different vital stains with different mechanisms of action. Fluorescein diacetate readily crosses the plasma membrane and is cleaved inside the cytosol of alive cells, releasing the brightly fluorescent fluorescein, that is retained inside alive cells. Fluorescence inside of protoplasts with filopods confirmed their viability (Fig. 2a). On the other hand, Evans blue does not penetrate the plasma membrane of live cells. While we observed dead protoplasts containing Evans blue staining, protoplasts with protrusions did not show Evans blue staining, again confirming their viability (Extended Data Fig. 4). These data show that protoplasts with protrusions are alive.

To further test protoplast viability, we analyzed their long-term behavior by following all three categories of protoplasts – smooth protoplasts, protoplasts with dots, and protoplasts with filopods -over a time course of 12 hours, taking an image once per hour (Fig. 2c, showing images every 4h). Analyzing the category filopods, we noticed that filopods can be retracted over time or, in very rare cases, emerge over time. The majority of protoplasts was viable over ≥12 h, and there was no major difference in viability between the three protoplast categories (Fig. 2b, Table S2). We thus conclude that protoplasts with protrusions are not more likely to die than smooth protoplasts and thus, that protrusions are not related to cell death.

Next, we hypothesized that the protrusions might be sticky material from the suspension, e.g. originating from pieces of membrane from disintegrated protoplast in the suspension. To test this hypothesis, we mixed plants with plasma membrane markers with two different fluorophores (*#*1: *p35S::GFP-LTi6B* and *#*2: *pUBQ10::LTi6B-TdTomato*) during the digestion and analyzed the suspension of mixed protoplasts (Fig. 2d). Across N = 3 independent experiments with a total of n = 574 alive protoplasts, we did not find a single green protoplast (#1) with magenta protrusions or a single magenta protoplast (#2) with green protrusions (Fig. 2d, Table S3). We thus conclude that the protrusions are part of the protoplasts and continuous with their plasma membrane.

### Composition of the protrusions reveals cortical identity

In theory, protrusions could be pure membrane extensions as they are encased by plasma membrane (Fig. 3a, Extended Data Fig. 5). Further investigating the content of the protrusions, we found that they can contain large structures such at ER and cytoskeletal elements (Fig. 3b-d, Extended Data Figs. 6,7). Note that not all protrusions contained ER or cytoskeletal elements. In particular, filopods containing microtubules were rare. However, the fact that protrusions can contain ER as well as microtubules shows that not only the plasma membrane of the protrusion is continuous with the protoplast, but also that the cytoplasm of the protoplast is continuous with the protrusions and confirmed again that protrusions appeared independent of the genetic marker.

Focusing on microtubules, we used two different marker lines: *p35S::GFP-TUA6,* which marks tubulin *α* and *p35S::GFP-MBD*, which marks microtubule bundles. The presence of microtubule arrays at the protoplast cortex could be verified in maximum intensity projections (Fig. 3c,d). In the *p35S::GFP-TUA6* marker line, microtubules are distinguishable from diffuse tubulin fluorescence by their straight shape and an increased GFP-TUA6 signal (Fig. 3c, Extended Data Fig. 7a). The fact that we additionally observed microtubules in filopods with the microtubule bundle marker *p35S::GFP-MBD* clearly established the presence of microtubule bundles in a subpopulation of protrusions (Fig. 3d, Extended Data Figure 7b). Furthermore, the filopods that contained microtubule signal, with either *p35S::GFP-TUA6* or *p35S::GFP-MBD* markers, appeared straighter than the average filopod marked by a plasma membrane marker. This increased persistence length suggests that protrusions can be mechanically reinforced by the presence of microtubular cytoskeleton (Fig. 3c,d; see also Fig. 4a, Extended Data Fig. 7).

Taken together, the cytoplasmic continuity between the protoplast and the protrusion further strengthens the point that the protrusions are not debris sticking to the protoplast membrane but indeed are continuous with the protoplasts. Furthermore, it reveals that the protrusions can have a cortical identity, begging the question of how they are formed, whether they can actively move and how they are regulated.

### Protrusion shape can be regulated by cytoskeleton

The previous analysis suggests that the protrusions can be diverse not only in shape (dots/filopods), but also in content. Can this be modulated? In order to modulate the shape of the protrusions, we aimed to stabilize the microtubule cytoskeleton. We focused our analysis on the *p35S::GFP-TUA6* line, in which microtubules are most likely to be slightly destabilized to start with^8^. We treated the plants with the microtubule stabilizing drug taxol for 1h prior to digestion and kept the protoplasts in taxol throughout the experiment. Control plants were treated with equivalent concentrations of DMSO.

Analyzing the GFP-TUA6 signal in the DMSO control condition (n = 300 protoplasts across N = 3 independent experiments), we observed diffuse GFP-TUA6 signal in 32 ± 5% of the protoplasts, as well as diffuse GFP-TUA6 signal with some microtubules in 46 ± 7% of protoplasts. Protoplasts with a strong microtubule signal were rare (8 ± 5%, Extended Data Fig. 8a, Table S4). The taxol-treated protoplasts (n = 298 protoplasts across N = 3 independent experiments) exhibited a significant stabilization of the microtubule cytoskeleton, as evidenced by a much higher frequency of protoplasts with strong microtubule signal (51 ± 13%), together with a similar level of diffuse GFP-TUA6 signal with some microtubule signal (30 ± 9%) and a strongly reduced fraction of protoplasts with diffuse GFP-TUA6 signal (6 ± 6%, Extended Data Fig. 8a, Table S4). These data confirm that taxol treatment stabilizes the microtubules in these conditions.

Next, we analyzed the protrusions containing a GFP-TUA6 signal. The overall number of protrusions (dots + filopods) containing any GFP-TUA6 signal was similar in taxol-treated protoplasts vs. DMSO control (17 ± 2% vs. 15 ± 3%, Table S4). Note that this is the fraction of protrusions marked with GFP-TUA6, which is much lower than the fraction of protrusions marked with a plasma membrane marker, see Fig. 1f. Analyzing the frequency of filopods containing a GFP-TUA6 signal, the DMSO data confirm that filopods containing a GFP-TUA6 signal is a rare event (6 ± 3%). The frequency of filopods containing a GFP-TUA6 signal was increased in the taxol-treated protoplasts (13 ± 1%, Table S4). In particular, the frequency of filopods containing a strong GFP-TUA6 signal and thus likely microtubules was increased in taxol-treated protoplasts: 10 ± 3 %, compared to only 1 ± 1 % in the DMSO control condition (Fig. 4 a). Note that these filopods had a remarkably straight shape (Fig. 4a), similar to what we observed in Fig. 3c,d, either sticking out from the round protoplast, or occasionally deforming the whole protoplast towards the filopods. Consequently, the fraction of protoplasts with tubulin containing dots was higher in the DMSO control compared to the taxol-treated protoplasts (9 ± 2% vs. 4 ± 1%). Thus, taxol seems to affect the distribution between tubulin-containing dots and filopods. Taken together, this shows that microtubule stabilization with taxol does not change the overall number of protrusions, but instead shifts the shape of the present protrusions towards microtubule-containing, straight filopods compared to more tubulin containing dots in the control condition, confirming that microtubule status can regulate the shape of the protrusions.

### Dynamics of the protrusions: passive movement

Next, we wanted to understand the dynamics of the protrusions. First, in timelapse movies, we observed that the filopods move (Fig. 4b, Extended Data Fig. 9a, Video S1). We thus wondered whether this movement is active or passive. In order to better understand how strong a passive movement to expect in our setup, we analyzed fluorescent polystyrene beads in the microwells in the same hyperosmotic buffer that we used for the protoplasts. Over time, the beads sank to the bottom of the wells and/or adsorbed to the side walls of the wells (Extended Data Fig. 9b). However, the fraction of beads that was free in the hyperosmotic buffer without contact to the walls of the wells stayed mobile and moved substantially, showing that there is considerable thermal motion in our system (Fig. 4b, Extended Data Fig. 9b). Comparing the movement of the beads to the movement of the filopods, we found that the motion is qualitatively similar (Fig. 4b, Extended Data Fig. 9a). This is consistent with the hypothesis that the observed motion of filopods is passive. While this does not completely exclude the possibility of an active movement, the most parsimonious conclusion is that of passive movement.

### Controlling the protrusions with osmolarity

Beyond cytoskeleton, a more obvious candidate regulator of the protrusions is osmolarity, as subjecting plant tissue to a hyperosmotic solution induced plasmolysis and reveals protrusions. To test the effect of osmotic changes on protoplasts, we subjected protoplasts (n = 97 protoplasts across N = 3 independent experiments) to cycles of hyperosmolarity (600 mM D-mannitol ∼ 720 mOsmol/kg) and close to iso-osmolarity (280 mM D-mannitol ∼ 340 Osmol/kg). We made use of the microwell system that allows to observe the exact same protoplasts at different osmolarities. We analyzed the behavior of the protoplasts during a decrease in osmolarity (i) from 600 mM to 280 mM D-mannitol (Fig. 4c, left), followed by an increase in osmolarity (ii) from 280 mM to 600 mM D-mannitol (Fig. 4c, right). After either transition, a third of the protoplasts did not change in terms of number and type of protrusions (white in Fig. 4c). A third of the protoplasts appeared sick or dead at any point during the experiment and were disqualified. A third of the protoplasts decreased the number or size of protrusions following the initial decrease in osmolarity (i), and a third of the protoplasts increased the number or size of protrusions during the following increase in osmolarity (ii) (green in Fig. 4c). We did not observe protoplasts increasing the number or size of protrusions following a decrease in osmolarity and rarely observed a decrease in the number/size of protrusions following an increase in osmolarity (red in Fig. 4c); Taking both transitions together, we further asked whether the same protoplasts both decreased their protrusions with a decrease in osmolarity and increased them with an increase in osmolarity (20 ± 6%), or whether they only did one (10 ± 7%) or the other (10 ± 9%). Figure 4d shows examples of each of these categories. Taken together, more than half of the alive protoplasts show at least either a decrease in protrusions in lower osmolarity or an increase in higher osmolarity. This demonstrates that protrusions are regulated by osmolarity.

To further challenge this conclusion, we observed protoplasts (n = 76 protoplasts across N = 3 independent experiments) subjected to increasing final concentrations of D-mannitol, increasing from 600 mM (∼720 mOsmol/kg) over 700 mM (∼850 mOsmol/kg) to 800 mM (∼980 mOsmol/kg) using the microwell system. As a control, we added hyperosmotic buffer solution of the same concentration, i.e. keeping the protoplasts at 600 mM D-mannitol (∼720 mOsmol/kg) during the whole experiment (n = 97 protoplasts across N = 3 independent experiments). We then analyzed whether the type, number and size of the protrusions stayed constant (white in Fig. 4e), increased (green in Fig. 4e, see Fig. 4f for examples), or decreased (red in Fig. 4e) with the addition of buffer, i.e. with increase in osmolarity (Fig. 4e, left), vs. keeping the same osmolarity in the control (Fig. 4e, right). Protoplasts that looked sick or dead at any point in the experiment were disqualified (grey in Fig. 4e). Comparing the increase of osmolarity to the control, we saw that the fraction of protoplasts that increase the type, number or size of protrusions with increasing osmolarity is much higher than those spontaneously increasing their protrusions in the control (37 ± 2% vs. 11 ± 2%). Conversely, the large majority of protoplasts in the control stayed unchanged (63 ± 4%), while this was only the case for 28 ± 2% of the protoplasts subjected to osmotic changes. We can further observe that changing the osmolarity is a stress for the protoplasts, as the fraction of dead protoplasts increased in the protoplasts experiencing an increase in osmolarity (33 ± 4% vs. 21 ± 2% in control). Taken together, these observations confirm that osmolarity can regulate the protrusions. Importantly, the population of protoplasts with increasing osmolarity showed 7/76 protoplasts across the N = 3 experiments that developed *de-novo* filopods from smooth protoplasts. This only occurred in 2/97 protoplasts in the control. This shows that protoplasts can develop protrusions *de novo* and that this behavior is increased by an increase in osmolarity.

Taken together, these data show that increasing osmolarity can not only induce extension of existing protrusions, but also lead to the formation of new protrusions. Thus, protoplasts are able to form *de novo* dots and filopods. This shows that protoplasts do not require a plasma membrane-cell wall attachment site to pull out the membrane during plasmolysis to form filopods.

### Filopods can aXach to the microwells

The discovery that a significant fraction of protoplasts makes protrusions opens the question of their biological functions in plant tissues. These functions are likely numerous, and some may echo the proposed role of Hechtian structures in mechanosensing ^6,9,10^ or in acting as a reservoir of membrane during plasmolysis^5,6^. Because we can see protrusions emerging d*e novo*, another, non-exclusive, function may relate to a role in actively reshaping the cell cortex in plants. This might include the local remodeling of the cell wall, the definition of new contact sites, or even the formation of secondary plasmodesmata. To support this claim, the formation of protrusions is not sufficient; protrusions must also attach to the cell wall. Accordingly, we would like to report our occasional but persistent observation that filopods can attach to the surface (Fig. 5a-c). This was most obvious when the tip of the filopod did not move over time (Fig. 5d). The rare formation of very long (>30 µm) filopods might be attributed to attachment to the surface (Extended Data Fig. 2). Sometimes, the tips of the attached filopods also showed small extensions, widening the surface area of the tip at the site of attachment (Fig. 5b,c). We hypothesize that these filopods might be forming *de novo* Hechtian attachment sites, engaging in a “search and capture” mechanism.

## Discussion

What does it take to form a protrusion? In animal cells, protrusions form *in vivo*, e.g. when the actin cytoskeleton polymerizes against the plasma membrane, pushing out lamellodopia and filopodia^11,12^. Tubules are also omnipresent on intracellular membranes in animal and plant cells, including the ER^13,14^, the Golgi apparatus^15,16^, as well as plastids^17^ in plants. Membrane tubes can be experimentally pulled out of membrane vesicles or cells by locally applying a force, e.g. using optical tweezers^18^ to pull a bead away from the membrane, by micropipette aspiration of a vesicle bound to a rigid surface^19^ or holding a vesicle by a micropipette and exerting a force by a fluid flow^20^, as well as by scaffolding the plasma membrane with membrane deforming proteins^21,22^, and by pulling on vesicles with molecular motors^23^. In plants, the emergence of Hechtian strands can be regarded as tube formation by pulling the membrane away from a contact: the membrane is attached to the cell wall at discrete attachment point, the Hechtian attachment sites, from where the Hechtian strands are pulled out as the protoplast retracts^6^. Beyond the Hechtian strands, protrusions can be induced artificially in protoplasts by overexpressing remorin, with the rationale that clusters of remorin would constrain the shape of the plasma membrane^24^.

Here we showed that plant protoplasts readily form protrusions that are continuous with the protoplast (Fig. 2d), can contain ER and cytoskeleton (Fig. 3), and are regulated in shape by the cytoskeleton (Fig. 4a) and in abundance by osmolarity (Fig. 4c-f). They further support the hypothesis that filopods might be forming *de novo* Hechtian attachment sites (Fig. 5). Our discovery thus suggests that protrusion formation is not a rare event, but rather a normal behavior observed in plants from different taxa. Our discovery therefore challenges the depiction of plant cells as “passive” and calls for a re-evaluation of our understanding of the role of the plant cell cortex.

Protrusions in plant protoplasts have been observed in specific contexts, notably when overexpressing remorin^24^, but why have protrusions not been observed earlier in normal protoplasts? Among many hypotheses, one can propose that (i) protrusions are small, dynamic structures and thus difficult to see in a sea of protoplasts, in particular in the absence of a bright plasma membrane marker (see Fig. 1b,e, Extended Data Fig. 2), (ii) “clean”, i.e. smooth protoplasts are usually selected as canonical protoplasts, and (iii) high centrifugation speed in the protoplast preparation protocol may damage protrusions. Our work might trigger re-analyses of previous results, which in turn may shed a new light on the role of these protrusions in specific contexts.

Are the protrusions Hechtian structures? The fact that new protrusions can emerge in hyper-osmotic conditions (Fig. 4f) suggests that protrusions are not simply remaining Hechtian strands from a plasmolyzed cell undergoing cell wall digestion. Yet, we will need to question the relation between filopods and Hechtian structures in further studies. The attachment of filopods to the surface of the microwell (Fig. 5a-d) suggests that protoplasts exhibit a competence to stick to surfaces through these protrusions, a behavior that is more animal-like than generally pictured.

The protrusions observed in plant cells can echo focal adhesions in animals, with the major difference that plant filopods appear passive. Yet, plant protrusions remain attractive candidates as mechanosensing hubs: the Hechtian attachment sites might bundle mechanical forces at discrete points on the plasma membrane; the protoplast protrusions might probe the environment for potential attachment sites. This also recalls the relation between adhesion and tension at the tissue scale, which is observed across kingdoms of life^25^. Our work suggests that this may very well be true at cellular scale, too. Conversely, the increasing appreciation of the role of osmolarity in animal cell and tissue morphogenesis^26,27^ might in turn echo the observed protrusions in plant protoplasts, further reflecting more universal behavior than anticipated.

## Methods

### Plant lines and growth conditions

#### Arabidopsis thaliana

For our analysis, we used the following stable transgenic *Arabidopsis thaliana* lines:

*pUBQ10::LTi6B-TdTomato*^28,29^, Col-0 background

*p35S::GFP-LTi6B*^30^, Col-2 background (NASC ID N84726)

*pUBQ10::DDRGK1-GFP*^31^, Col-0 background

*p35S::GFP-TUA6*^32^, Col-0 background (NASC ID N6551)

*p35S::GFP-MBD*^33,34^, WS-4 background

Arabidopsis seeds were sterilized in 70% ethanol for 10 minutes, dried on filter paper and transferred to ½ MS plates (Murashige & Skoog Basalt salt mixture, Duchefa), 3 mM MES hydrate (Sigma), 0.8% plant agar (Duchefa), pH adjusted to 5.7 using KOH, without sugar or vitamins). Plates were cold-stratified at 4°C in the dark for 2-3 days and then grown in long day plant growth chambers (16h light/ 8h dark, 20°C) for 7 days unless otherwise noted.

#### Physcomitrium patens

Homogeneous protonemal cultures (Gransden) were grown for 4-7 days under long-day conditions (16h light/ 8h dark) on BCDAT media plates (250mg/l MgSO_4_.7H_2_O, 250mg/l KH_2_PO_4_ (pH6.5), 1010mg/l KNO_3_, 12.5mg/l FeSO_4_7H_2_O, 0.001% Trace Element Solution (0.614mg/l H_3_BO_3_, 0.055mg/l AlK(SO_4_)_2_.12H_2_O, 0.055mg/l CuSO_4_.5H_2_O, 0.028mg/l KBr, 0.028mg/l LiCl, 0.389mg/l MnCl_2_.4H_2_O, 0.055mg/l CoCl_2_.6H_2_O, 0.055mg/l ZnSO_4_.7H_2_O, 0.028mg/l KI and 0.028mg/l SnCl_2_.2H_2_O), 0.92 g/l C_4_H_12_N_2_O_6_, 1mM CaCl_2_ and 8g/l plant agar) overlaid with sterile cellophane discs.

#### Zea mays

Seedlings were grown for 6 days in a Petri dish on wet paper in the dark at 20°C.

### Buffer solutions

Hyperosmotic buffer solution: 600 mM D-mannitol, 1 mM CaCl_2_, 1 mM MgCl_2_, 10 mM MES in milliQ water, pH adjusted to 5.5 with KOH. The buffer was autoclaved and used at room temperature. In Fig. 4c-f, the D-mannitol concentration of the buffer was varied as stated in the figure. Osmolarity measurements reflect the osmolarity of the final buffer solution including the stated D-mannitol concentration and were performed on a cryoscopic osmometer (Osmomat030, Gonotec). In Fig. 2d and Extended Data Fig. 1b, D-sorbitol was used instead of d-mannitol.

Digestion buffer: Hyperosmotic buffer with 600 mM D-mannitol as described above, including additionally 17 mg/ml Cellulysine (Merck Millipore), 17 mg/ml Cellulase RS (Duchefa), 0.4 mg/ml Pectolyase Y23 (Duchefa), 3.5 mg/ml BSA (Sigma), 1 mM ascorbic acid. The digestion buffer was filtered through a 0.45 µm filter and frozen in 1.1 ml aliquots in 2 ml round-bottom Eppendorf tubes at -20°C. In Fig. 2d and Extended Data Fig. 1b, D-sorbitol was used instead of d-mannitol.

### Plasmolysis of Arabidopsis hypocotyl

7d old *pUBQ10::LTi6B-TdTomato* plants (not cold stratified) were exposed to hyperosmotic buffer solution for 50 min in a 6-well plate, then transferred to a glass slide with a custom-made vacuum grease (Molykote, Dupont) chamber containing 500 µl hyperosmotic buffer solution, covered with a cover slip and imaged on a LSM980 upright confocal microscope (Carl Zeiss Microscopy GmbH, Jena), controlled by the Zen blue 3.6 software, with Objective W Plan-Apochromat 20x/1.0 (water immersion), and the 561 nm laser line, a GaAsP PMT detector for the fluorescent signal, and a multialkali PMT detector for the transmitted light signal.

### Preparation of protoplasts

#### Arabidopsis

Preparation of Arabidopsis protoplast is adapted from the description in Colin *et al*._7_: Ca. 30 7-day old Arabidopsis seedlings were added to a de-frosted 1.1 ml aliquot of the digestion buffer and agitated in a 2 ml round-bottom Eppendorf tube on a rotary shaker at 15 rpm in a plant growth chamber at 20°C for 45min. The plants were carefully pipetted up and down three times using a p1000 micropipette with a cut tip and centrifuged at 100 g for 4min. The supernatant including the plants was removed by inverting the tube, the pellet was resuspended in 1 ml hyperosmotic buffer solution by carefully inverting the tube and centrifuged at 100 g for 4min. The supernatant was removed by inverting the tube and the pellet was carefully resuspended in the remaining liquid, ca. 100 µl hyperosmotic buffer solution, by flicking the tube.

#### Physcomitrium patens

Protonemal tissue was incubated for 1h in a sterile solution of 10 mg/ml Driselase in 8% w/v D-mannitol solution. The resulting mixture was filtered twice through a sterile 70 µm nylon filter. Protoplasts were sedimented by spinning at 70 g for 5 minutes, and rinsed three times in 8% w/v D-mannitol solution before observation. Protoplasts were imaged in 8% w/v D-mannitol solution containing FM4-64 at a final concentration of 2 µg/ml in a NOA container. Filopods were observed in N = 3 independent experiment.

#### Maize

Maize leaves and roots were digested in the same digestion buffer as the Arabidopsis plants for 45 min at 15 rpm in an Eppendorf tube. After digestion, the protoplasts were filtered through a 70 µm filter and 1 ml of the hyperosmotic buffer solution was added to wash the filter and the protoplasts. After sedimenting the protoplasts for ca. 1h, the supernatant was removed with a pipette and the pellet was washed with 1 ml hyperosmotic buffer solution. After another hour of sedimentation, the supernatant was removed and the pellet was resuspended in a final volume of ca. 50 µl of hyperosmotic buffer solution. The protoplasts were stained with FM4-64 at a final concentration of 2 µg/ml and imaged in a NOA73 container. Fig. 1h shows a representative image of N = 2 experimental repeats with this protocol, we obtained similar results with Maize leaves from 12 days old plants grown on soil in the dark, centrifuged at 100 g for the washing steps (N = 1 experiment).

### NOA73 container or microwells for imaging the protoplasts

A custom imaging support for the protoplast was generated by making NOA73 (Norland Optical Adhesive 73, an optically clear, liquid, UV-curable adhesive, Norland Optical Adhesive 73, Norland Products Inc.) containers or microwells in an Ibidi glass bottom dish (µ-Dish, 35 mm high, Ibidi GmbH, Gräfelfing). A drop of NOA73 was added to the glass bottom of an Ibidi dish and a PDMS (Polydimethylsiloxane, Sylgard 184, Dow Inc., Midland) stencil with micropillars of 15 x 20 x 21 µm (to generate NOA73 microwells, as in^7^) or a piece of PDMS (4 x 4 x 2 mm, to generate a NOA73 container) was pressed into the NOA73. The NOA73 was polymerized using a UV LED illuminator (UV-KUB 2, Kloé SA, Montpellier) at 100% for 3 min. The Ibidi dish was filled with milliQ water and the PDMS stencil was removed in the water. The microwells/container were washed 2 x 5 min with hyperosmotic buffer solution.

### Protoplast Imaging

A drop of protoplast suspension (ca. 30 µl) was added carefully to the top of the custom-made imaging support and left to sediment for 15 min. 3 ml hyperosmotic buffer solution was added and the Ibidi dish was closed. Note that for experiments reported in Fig. 4e,f, 1 ml hyperosmotic buffer solution was added; for Fig. 4c-f the dish was left open during the imaging to allow for buffer replacement/addition. The protoplasts were imaged on an inverted confocal LSM 980 microscope (Carl Zeiss Microscopy GmbH, Jena) controlled by the Zen blue 3.6 software, Objective C-Apochromat 40x/1.2 W (water immersion), using the 488 nm laser line, the 561 nm laser line, the 639 nm laser line (for Evans blue experiments, only), GaAsP PMT detectors for the fluorescent signal, and a multialkali PMT detector for the transmitted light signal. For multi-channel imaging, two GaAsP detectors were used, allowing to acquire all channels at the same time to prevent shifts between channels due to movement of the protrusions.

### Quantification of protoplasts

All experiments were analyzed manually using Fiji^35^. For all quantifications, the whole z-stack was analyzed, even though we sometimes display the maximum intensity projection for ease of visualization. Note that often protrusions are present close to the poles of the protoplasts that are not visible in a maximum intensity projection. The relative occurrence of observed outcome (e.g. filopods, microtubule signal, etc.) were averaged across experimental repeats.

For the protoplast statistics presented in Fig. 1f, protoplasts were contained in a NOA73 container. The lower rim of the container was systematically imaged, without selecting fields of views. The alive protoplasts in the first 10-20 fields of views were manually classified in smooth, dots, filopods, using the Cell Counter plugin (https://github.com/fiji/Cell_Counter/blob/Cell_Counter3.0.0/src/main/java/sc/fiji/cellCounter/Cell_Counter.java) in Fiji and counted. N = 3 experimental repeats with a total of 218, 470, 234 alive protoplasts per repeat, respectively. The relative occurrence of the three categories across the three repeats was averaged and the standard deviation was calculated.

### FDA staining

FDA (fluorescein diacetate, Sigma) stock solution: 5 g/l in acetone. FDA staining solution: 50 µg/ml in hyperosmotic buffer solution. Protoplasts were prepared as described above and imaged in 3 ml hyperosmotic buffer solution, recording the positions (Zen software). 150 µl of the FDA staining solution were added into the Ibidi dish, resulting in a final FDA concentration of 2.5 µg/ml. The same protoplasts were imaged again 10-25min after adding the staining solution. N = 3 independent experiments.

### Evans blue staining

Protoplasts were prepared as described above with 2 differences: The centrifugation speed was 1000 g, an additional washing/centrifugation step was added. After the first centrifugation step, Evans blue (Sigma) staining solution was added at a final concentration of 0.1 g/l (stock solution 10 g/l in water). Protoplasts were washed in 1 ml hyperosmotic buffer and then imaged. Evans blue is fluorescent when excited at 639 nm. Presence or absence of blue staining was also verified through eye-pieces. No Evans blue was detected inside of protoplast with filopods in N = 3 independent experiments with protoplasts obtained from 4-day old (2 experiments) and 6-day old plants (1 experiment), respectively.

### Overnight time course

For the overnight time course, the protoplasts were imaged in microwells in an Ibidi dish containing 3 ml hyperosmotic buffer solution with the lid closed to minimize evaporation during the night. Immersion oil (Immersol W (2010), Carl Zeiss Microscopy GmbH, Jena) with a refractive index of water (n = 1.334) was used to avoid evaporation of immersion medium during the night. The same protoplasts were imaged once every hour using the position list function (Zen software). Viability was assessed based on fluorescence of the plasma membrane marker, as well as the morphology of the protoplast in the transmitted light image. N = 3 experimental repeats with 27, 35, 35 protoplasts, respectively. For the first repeat, the protoplasts were not centrifuged during the protoplast preparation. Instead, the plants were carefully pipetted up and down after 45 min of digestion and 1 ml hyperosmotic buffer was added to stop the digestion. The sample was left to sediment for 30 min, 2 ml of digestion buffer was removed with a pipette, protoplasts were washed with 1 ml hyperosmotic buffer, left to sediment for 30 min, 1.3 ml buffer was removed with a pipette and the protoplasts resuspended in the remaining hyperosmotic buffer solution. For the other two repeats, protoplast preparation as described above.

### Mixed protoplasts

15 plants each of *pUBQ10::LTi6B-TdTomato* and *p35S:GFP:LTi6B*, each 8d old, 2d cold stratified were jointly digested in a 2 ml round-bottom Eppendorf tube with digestion buffer containing 0.6 M D-sorbitol instead of D-mannitol and imaged in 3 ml hyperosmotic buffer solution containing 0.6 M D-sorbitol instead of D-mannitol. For imaging, protoplasts were contained in a NOA73 container. The tile region module (Zen software) was used to systematically acquire the whole lower rim of the NOA73 container. TdTomato and GFP were excited at same time and the fluorescence signal was separated by detection filters, acquiring the two signals simultaneously on two different GaAsP detectors to avoid movement of filopods in between frames. N = 3 experimental repeats with 324, 159, and 91 alive protoplasts each.

### FM4-64 staining

For the Arabidopsis experiments (Fig. 3b-d, Extended Data Figs. 6,7), the FM4-64 was added directly to the Ibidi dish containing 3 ml of hyperosmotic buffer solution and the protoplasts to a final concentration of 0.5 µg/ml and imaged 10 min after addition of the dye.

### Taxol treatment

Ca. 30 plants were incubated for 1h – 1h15min in 1 ml ½ MS medium (0.5x Murashige & Skoog Basalt salt mixture (Duchefa), 3 mM MES hydrate (Sigma), pH adjusted to 5.7 using KOH) containing 2 µl of taxol (paclitaxel, Sigma) stock solution (20 mM in DMSO), i.e. final concentration 40 µM, or 2 µl DMSO as a control on a turning plate at 15 rpm in the growth chamber at 20°C. Then, plants were transferred to a 2 ml Eppendorf tube containing 1.1 ml digestion solution with 1.1 µl taxol stock solution, i.e a final concentration of 20 mM, or 1.1 µl DMSO (control) and digested in a turning plate at 15 rpm in the growth chamber at 20°C for 45-55 min. Plants were pipetted up and down carefully, the sample was centrifuged at 100 g for 4 min and washed with 1 ml hyperosmotic buffer containing taxol at a final concentration of 20 mM or equivalent volume of DMSO (control). Protoplasts were centrifuged at 100 g for 4min, the supernatant was decanted and the pellet was carefully resuspended in the remaining hyperosmotic buffer solution (ca. 100 µl). A drop of protoplast suspension was transferred to microwells on an Ibidi dish, left to sediment for 15-20min and 3 ml hyperosmotic buffer containing taxol at a final concentration of 20 µM, or an equivalent volume of DMSO was added. 100 protoplasts per experimental repeat and per treatment vs. control were selected for imaging based on their strong fluorescence without selecting a specific protoplast category (smooth/dots/filopods) and z-stacks were acquired automatically using the position list option (Zen software). (Difference for repeat 1: 6-day old plants without cold stratification, 98 protoplasts in taxol condition).

### Movement assay

Protoplasts (*pUBQ10::LTi6B-TdTomato*) were imaged in NOA73 microwells in 3 ml hyperosmotic buffer solution in a closed Ibidi dish at maximum acquisition speed in order to capture the maximum of movement possible (7x zoom, bidirectional imaging, no averaging, maximum imaging speed (frame time: 103ms), timelapse of a single confocal plane). Multifluorescent polystyrene beads (diameter: 1 µm, Fluoresbrite, 24062, Polysciences Inc.) were imaged in NOA73 microwells prepared in the same way as for the protoplasts in 3 ml hyperosmotic buffer with the same imaging parameters. To visualize the movement, 20 consecutive frames of the fluorescent channel of the time lapse were color-coded by time and superposed using the Temporal Color Coder macro (https://imagej.net/ij/macros/Temporal-Color_Code.txt) in Fiji to visualize the movement. The final images were smoothed with a Gaussian filter, σ = 1. The same visualization (without Gaussian smoothing) was used for the representation of filopods movement in Fig. 5d. Note that protoplasts presented in Fig. 5b-d were obtained from 6-day old plants without cold stratification, using 1000 g for the centrifugation steps. A drop of 30 µl protoplast suspension was left to sink into the microwells for 1h and then imaged.

### Cycles of osmolarity

Protoplasts (*pUBQ10::LTi6B-TdTomato*) were contained in NOA73 microwells in an open Ibidi dish in 3 ml hyperosmotic buffer solution (600 mM D-mannitol). The dish was scanned for brightly fluorescent protoplasts of all categories (smooth, dots, filopods) and a position list (Zen software) was generated. z-stacks of all protoplasts on the list were acquired, 2.5 ml hyperosmotic buffer solution were carefully removed with a pipette without moving the dish and 3 ml hyperosmotic buffer solution with 280 mM D-mannitol were added. The focus of all positions in the position list was checked and updated if necessary and the whole list was re-imaged 5min after changing the buffer solution. Afterwards, 3 ml buffer solution were carefully removed using a pipette and replaced by 3 ml hyperosmotic buffer solution (600 mM D-mannitol). The focus was again verified and imaging was re-started 5-10min after changing the buffer solution. N = 3 experimental repeats with 30, 31, and 36 protoplasts, respectively.

### Increase in osmolarity

Protoplasts (*pUBQ10::LTi6B-TdTomato*) were contained in NOA73 microwells in an open Ibidi dish in 1 ml hyperosmotic buffer solution (600 mM D-mannitol). Brightly fluorescent protoplasts of all three categories (smooth, dots, filopods) were selected and saved in a position list (Zen software). z-stacks of the position list were imaged, and 1 ml hyperosmotic buffer solution with 800 mM D-mannitol was added, leading to a final concentration of 700 mM D-mannitol. The position list was re-imaged 5min after adding the buffer solution (immediately in one of the repeats). Afterwards, 1 ml of hyperosmotic buffer solution with 1 M D-mannitol was added, leading to a final concentration of 800 mM D-mannitol and the position list was imaged again after 5-10min (immediately in one of the repeats).

N = 3 experimental repeats quantified with 20, 25, and 31 protoplasts, respectively. 4^th^ experimental repeat was not included in quantification, as in this repeat >50% of protoplasts died during the imaging. As a control, the same experimental set-up was used, but 1 ml hyperosmotic buffer solution with 600 mM D-mannitol was added at both occasions, keeping the osmolarity constant across the experiment. N = 3 experimental repeats with 31, 30, and 36 protoplasts, respectively.

## Data availability

All raw images analyzed in this study have been deposited with unrestricted access at zenodo.org, DOI: 10.5281/zenodo.14107989). Other materials or data are available on demand and/or in the main text and supplementary material.

## Acknowledgements

We thank the MechanoDevo team and RDP lab for insightful discussions and Yvon Jaillais’ team and Gwyneth Ingram’s team for kindly sharing seeds. Protrusions were observed initially during a stay at the Mechanobiology Institute (Singapore), with Tim Saunders and Virgile Saunders (2019). We also thank Naomi Nakayama, who also observed similar structures in her team (Imperial College, London), consistent with our observations. This work was funded by the Deutsche Forschungsgemeinschaft (DFG, German Research Foundation) – project number 521501033 to J.D. and the European Research Council (ERC-2021-AdG-101019515 “Musix” to O.H.).

## Author contribution

M.M. and C.L. contributed equally. Conceptualization: J.D., O.H. Methodology: J.D, M.M, C.L., Z. N-V., O.H. Funding acquisition: J.D., O.H. Project administration: O.H. Supervision: J.D., O.H. Writing – original draft: J.D. Writing – review & editing: J.D, C.L., Z. N-V., M.M, O.H

---

**Extended Data Fig. 1.**
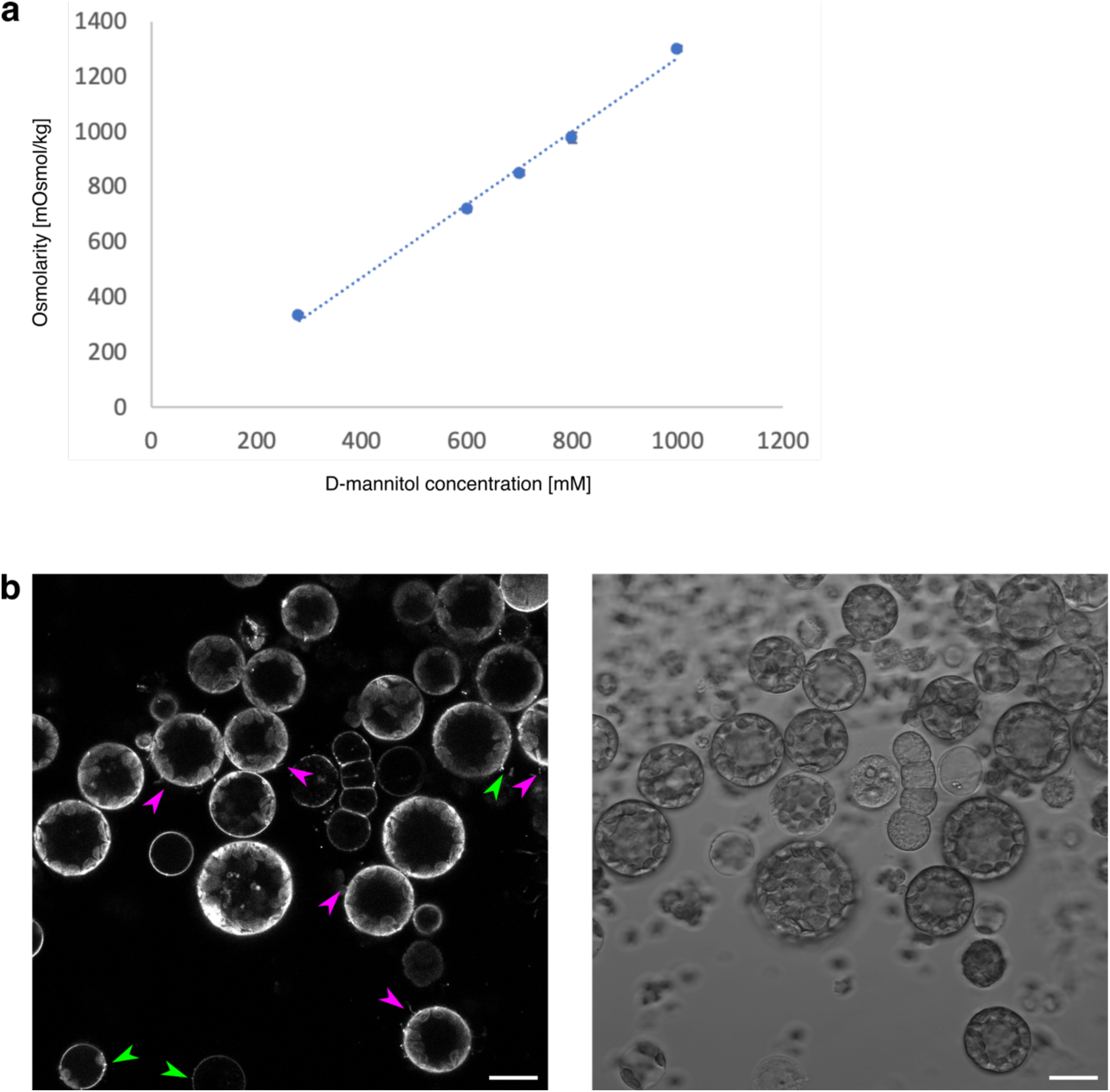
Osmolarity of hyperosmotic buffer solution and protoplasts in hyperosmotic buffer containing D-sorbitol instead of D-mannitol. **a**, Osmolarity of hyperosmotic buffer solution. Measured osmolarity of the hyperosmotic buffer of different D-mannitol concentrations. Dots show average of three (700, 800, 1000 mM) or six (280 mM, 600 mM) technical repeats of measurement. Error bars show standard deviation but are smaller than the markers. Dotted line: trend line showing linear fit. R^2^ = 0.9942. **b**, Overview image of protoplasts digested and imaged in hyperosmotic buffer solution containing D-sorbitol instead of D-mannitol. Left: *pUBQ10::LTi6B-TdTomato* (plasma membrane marker). Single confocal plane. Protoplasts with dots are marked with green arrow heads, protoplasts with filopods are marked with magenta arrow heads. Right: transmitted light image. Scale bars: 20 µm. LUT of both images was set to 0-150.

**Extended Data Fig. 2.**
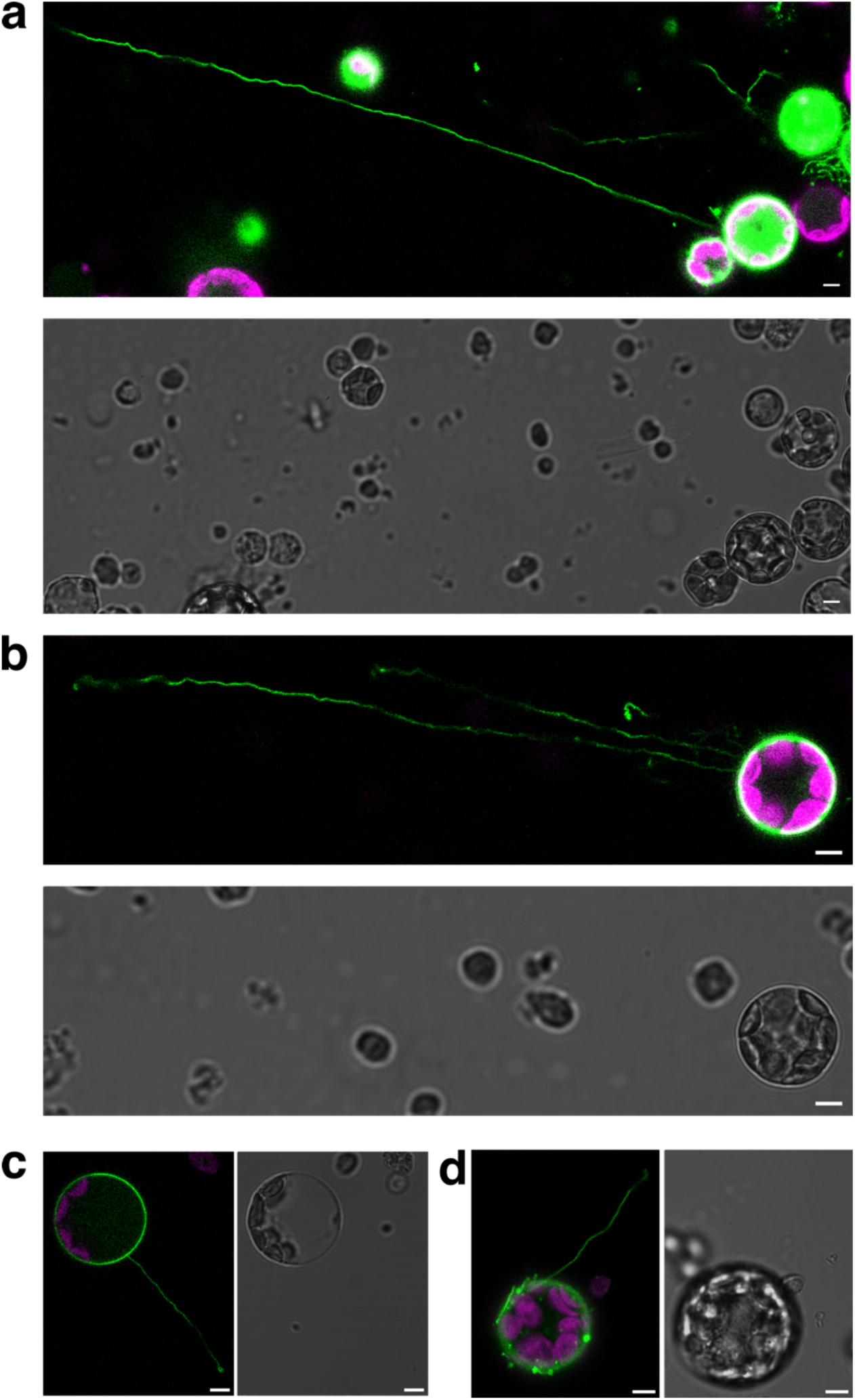
Examples of protoplasts with very long filopods. **a-d**, Protoplasts with a plasma membrane marker (*pUBQ10::LTi6B-TdTomato*). Upper/Left: overlay of membrane signal (pseudo-colored in green) and chloroplast autofluorescence (pseudo-colored in magenta), single confocal plane. Lower/Right: transmitted light. Protrusion lengths: ca. 210 µm (**a**), ca. 125 µm (**b**), ca. 40 µm (**c**), ca. 35 µm (**d**). **a**, **b**, Protoplasts from 8-day-old plants, observation of a 20 µl drop of protoplast suspension in a NOA73 container without adding additional hyperosmotic buffer solution. **b**, LUT transmitted light: 0-120. **c**, **d**, Observation between glass and cover slip. **c**, protoplasts from 7-days old plants, LUT green channel and transmitted light: 0-150. **d**, protoplasts from 6-day old plants. LUT transmitted light: 0-120. All scale bars: 5 µm

**Extended Data Fig. 3.**
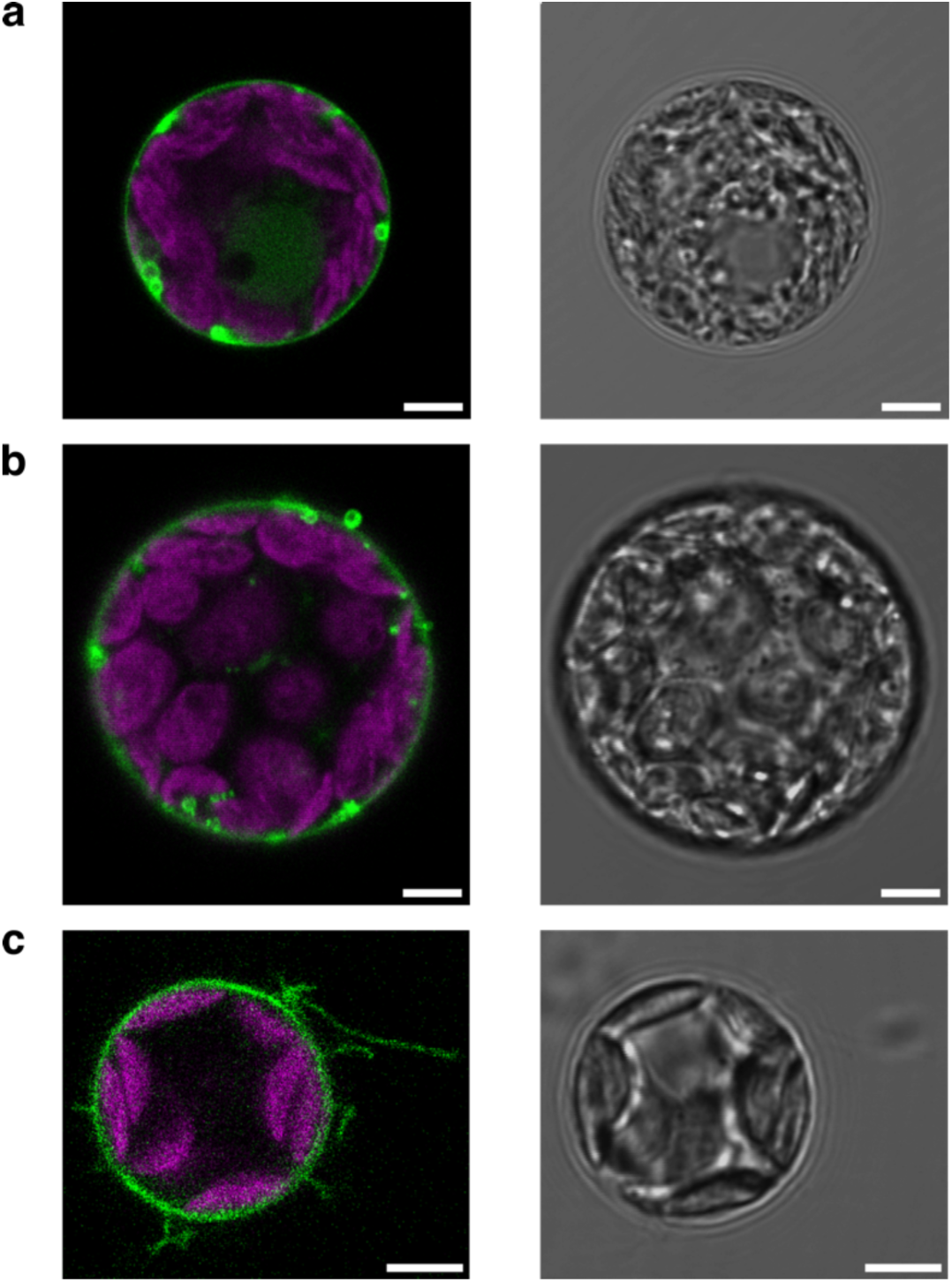
Protoplast on glass slide. Protoplasts with a plasma membrane marker (*pUBQ10::LTi6B-TdTomato*) in hyperosmotic buffer solution between a glass slide and a cover slip. Left: membrane marker signal pseudo-colored in green, overlayed with chloroplast autofluorescence signal pseudo-colored in magenta. Right: transmitted light. **a**, Smooth protoplasts. **b**, Dots. **c**, Filopods. Scale bars: 5 µm. **a**, **b** protoplasts from 6-day old plants, LUT transmitted light: 0-120. **c**, protoplast form 7-day old plant, LUT green: 0-150, LUT transmitted light: 0-60.

**Extended Data Fig. 4.**
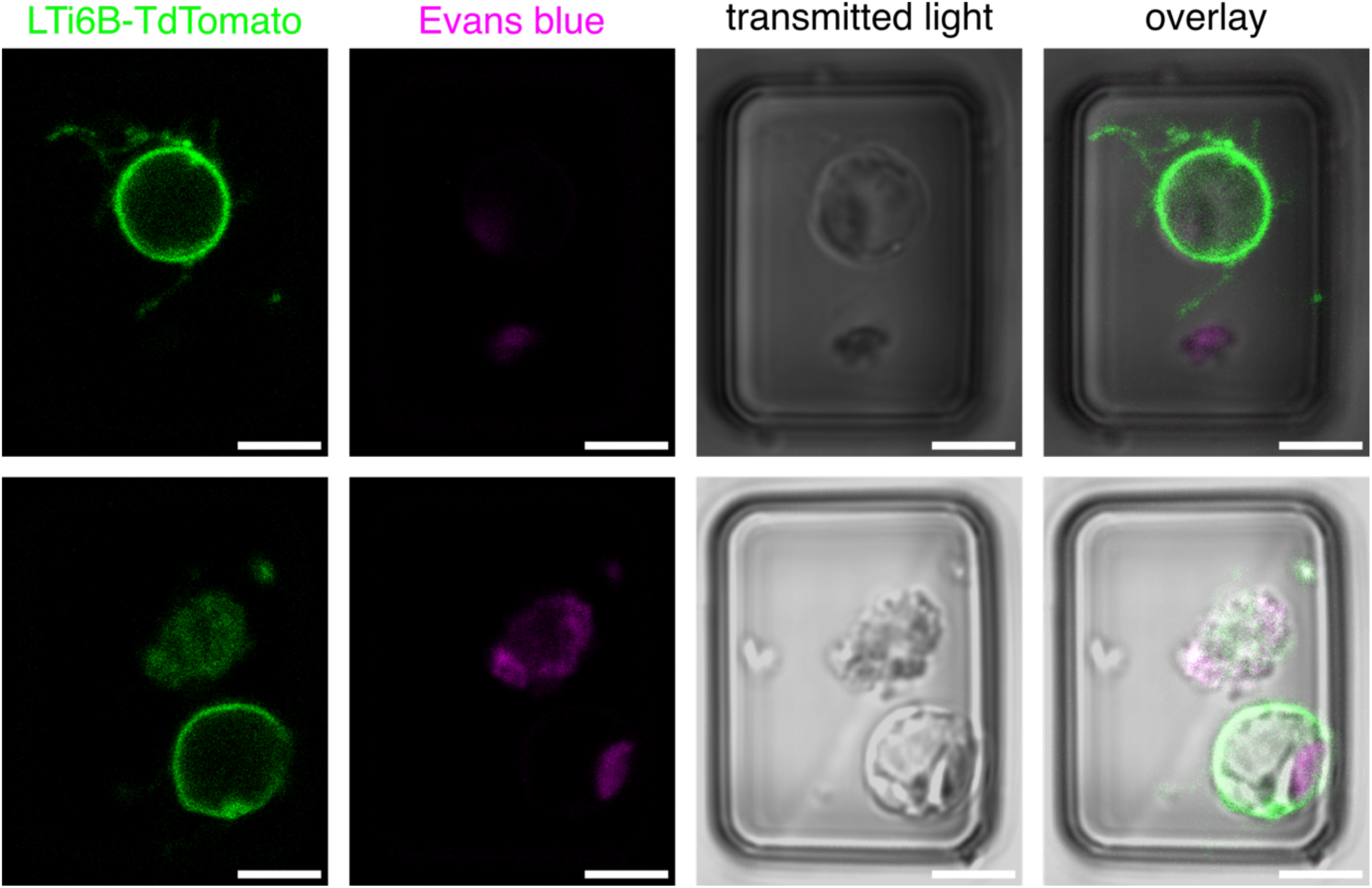
Protoplast viability test with Evans blue staining. From left to right: *pUBQ10::LTi6B-TdTomato* signal (pseudo-colored in green), Evans blue signal (pseudo-colored in magenta), transmitted light signal, overlay. Upper row: Example of protoplast with filopods without Evans blue signal, i.e. alive. Lower row: Example of smooth protoplast without Evans blue signal, i.e. alive, next to debris with Evans blue signal, i.e. dead (magenta signal inside of protoplast is a chloroplast as visible by the shape). Fluorescent images show single confocal plane. Scale bars 5 µm. We did not detect Evans blue signal in protoplasts with filopods across N = 3 independent experiments.

**Extended Data Fig. 5.**
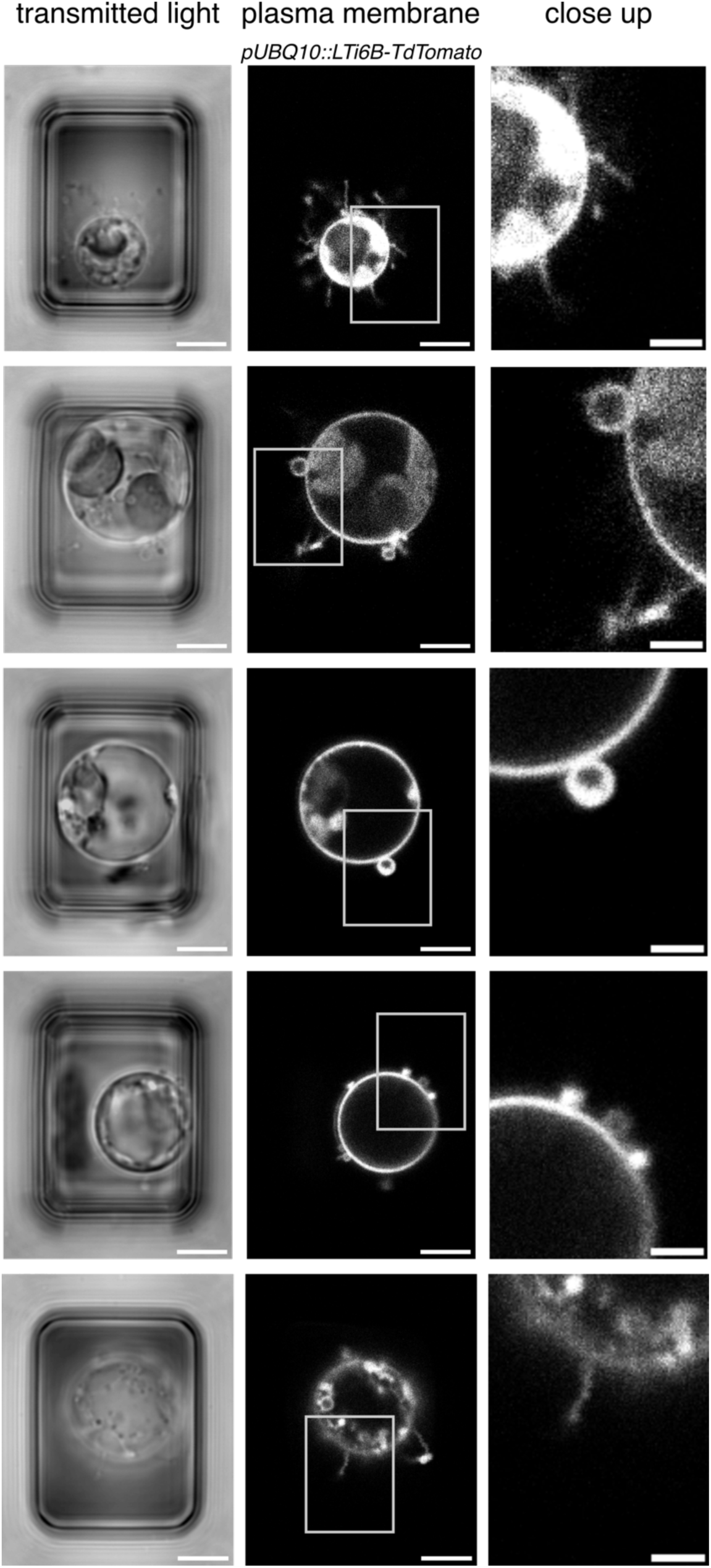
Additional examples of protrusions: Plasma membrane marker. *pUBQ10::LTi6B-TdTomato* (plasma membrane marker). Transmitted light image (left), single confocal plane (middle), close up as marked by box in the middle (right). Scale bars: 5 µm, close up: 2 µm.

**Extended Data Fig. 6.**
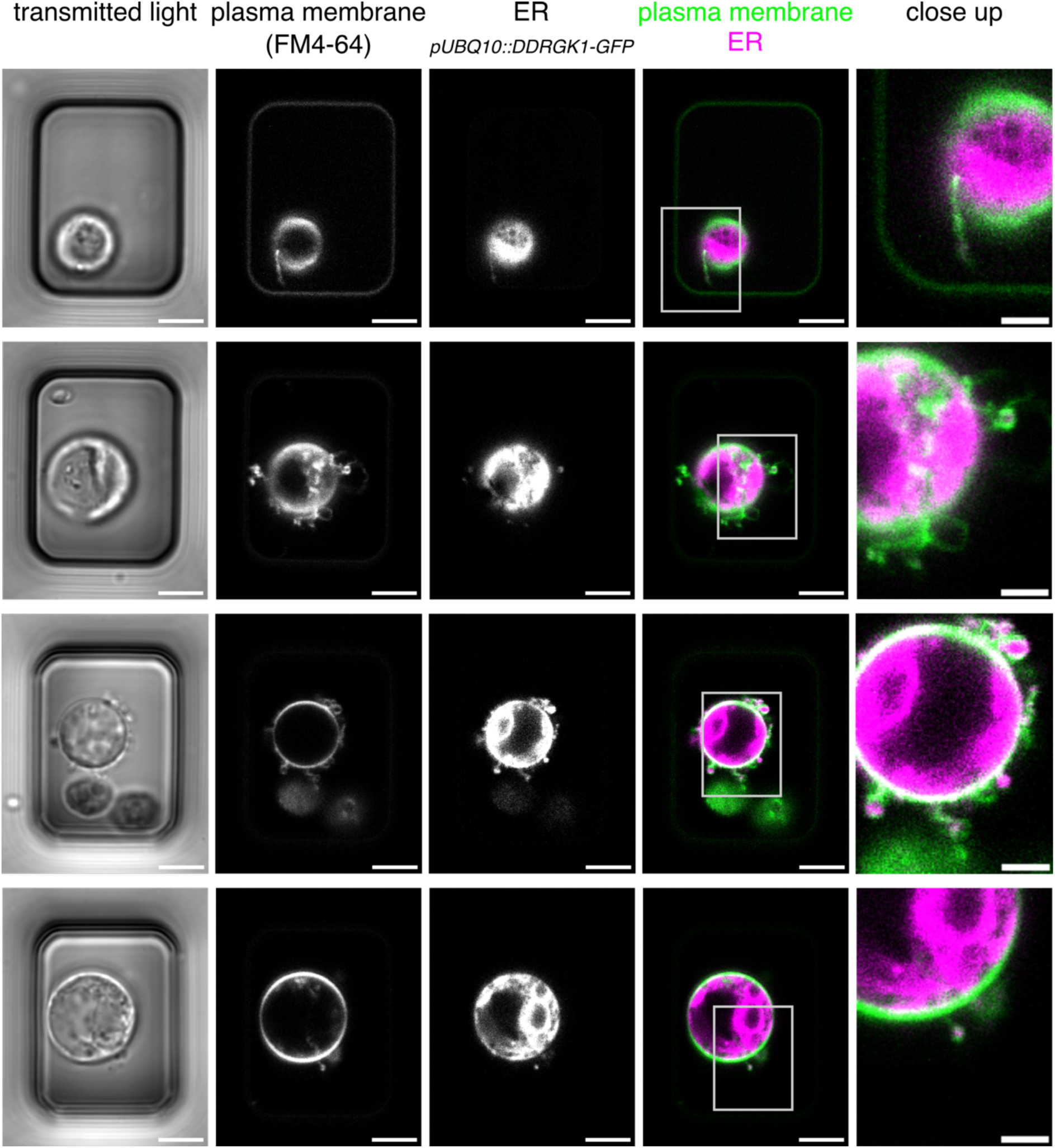
Additional examples of protrusions: ER membrane marker. *pUBQ10::DDRGK1-GFP* (ER membrane marker). Close ups as specified by box in precedent column. Fluorescent images show single confocal planes. Scale bars: 5 µm, close ups: 2 µm.

**Extended Data Fig. 7.**
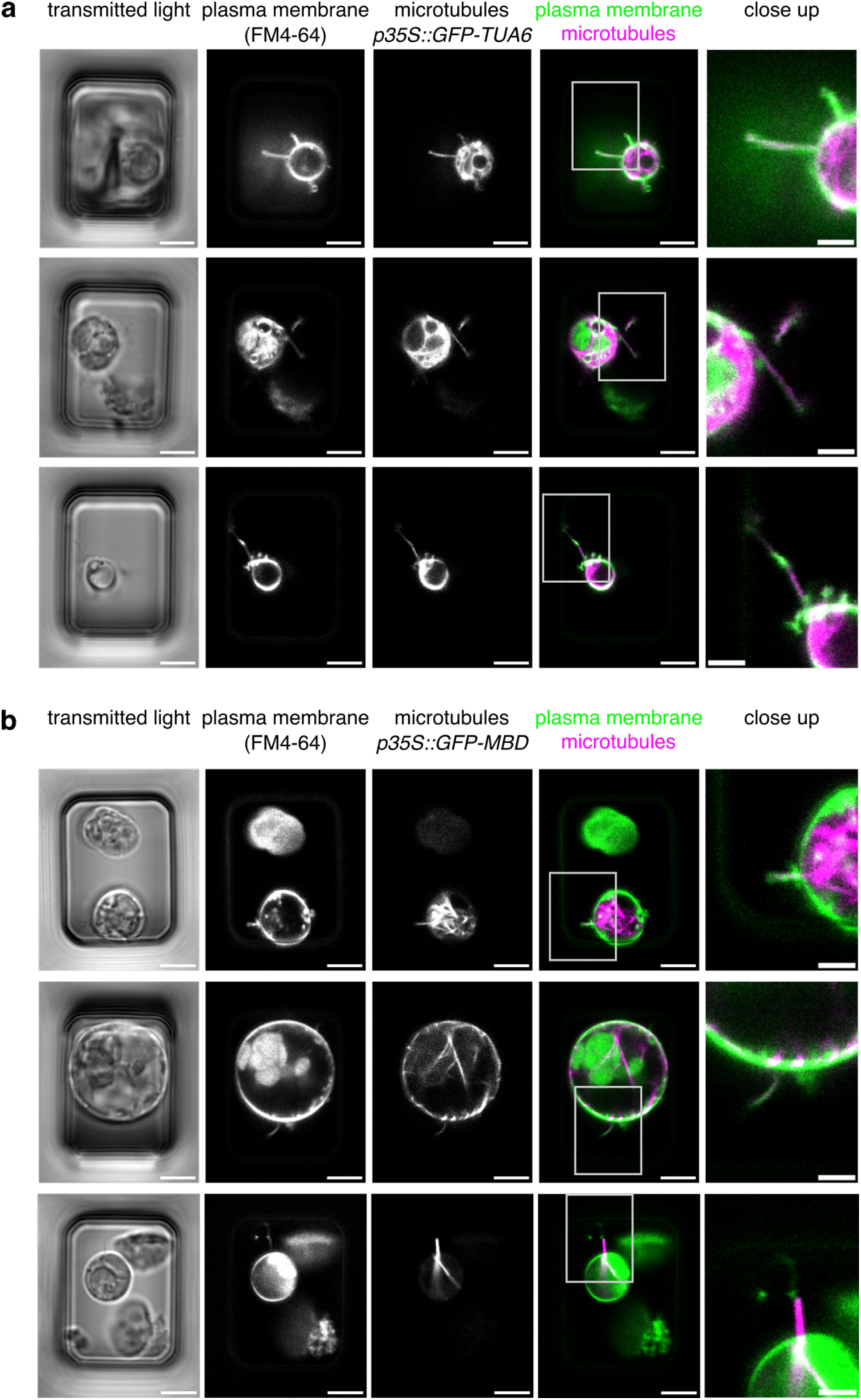
Additional examples of protrusions: Microtubule markers. **a**, *p35S::GFP-TUA6* (tubulin marker). **b**, *p35S::GFP-MBD* (microtubule bundle marker). Close ups as specified by box in precedent column. Fluorescent images show single confocal planes. Scale bars: 5 µm, close ups: 2 µm.

**Extended Data Fig. 8.**
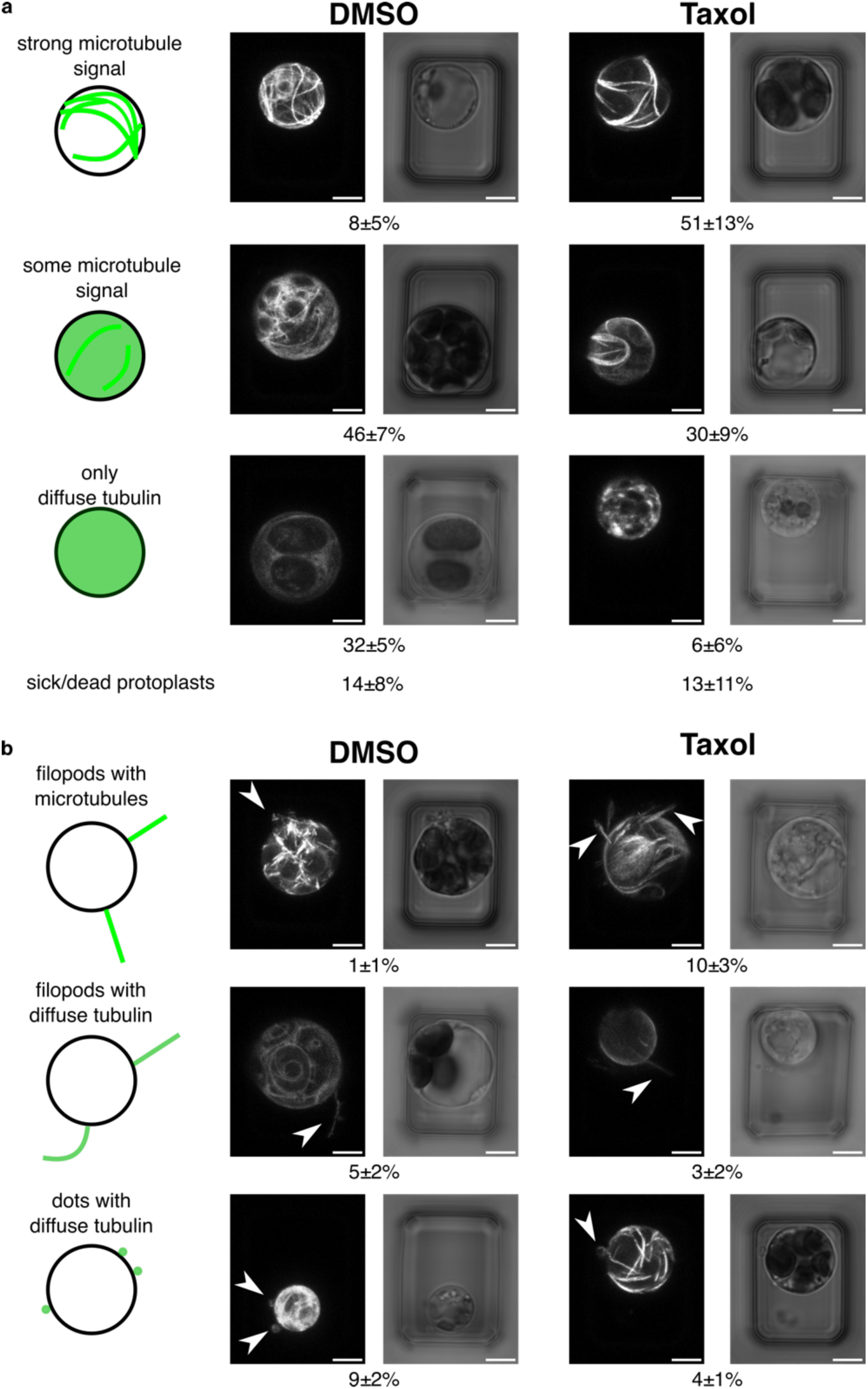
Taxol stabilizes microtubules and changes the shape of tubulin-containing protrusions towards filopods. **a**, Example images (maximum intensity projection of tubulin marker (*p35S::GFP-TUA6*) and transmitted light), and average fractions ± standard deviation of protoplasts with strong GFP-TUA6 microtubule signal (top), some GFP-TUA6 microtubule signal and some diffuse GFP-TUA6 signal (middle), and diffuse GFP-TUA6 signal (bottom). The numbers for dead/sick protoplasts are stated in the last row. **b**, Fractions of all protoplasts that show GFP-TUA6 signal in their protrusions classified based on protrusion shape and intensity of GFP-TUA6 signal as indicated on the left. Maximum intensity projection of GFP-TUA6 signal (*p35S::GFP-TUA6*) and transmitted light images. Arrow heads point towards protrusions. Scale bars: 5 µm. Percentages state average fraction ± standard deviation of the respective protrusions over n = 298 protoplasts across N = 3 independent experiments.

**Extended Data Fig. 9.**
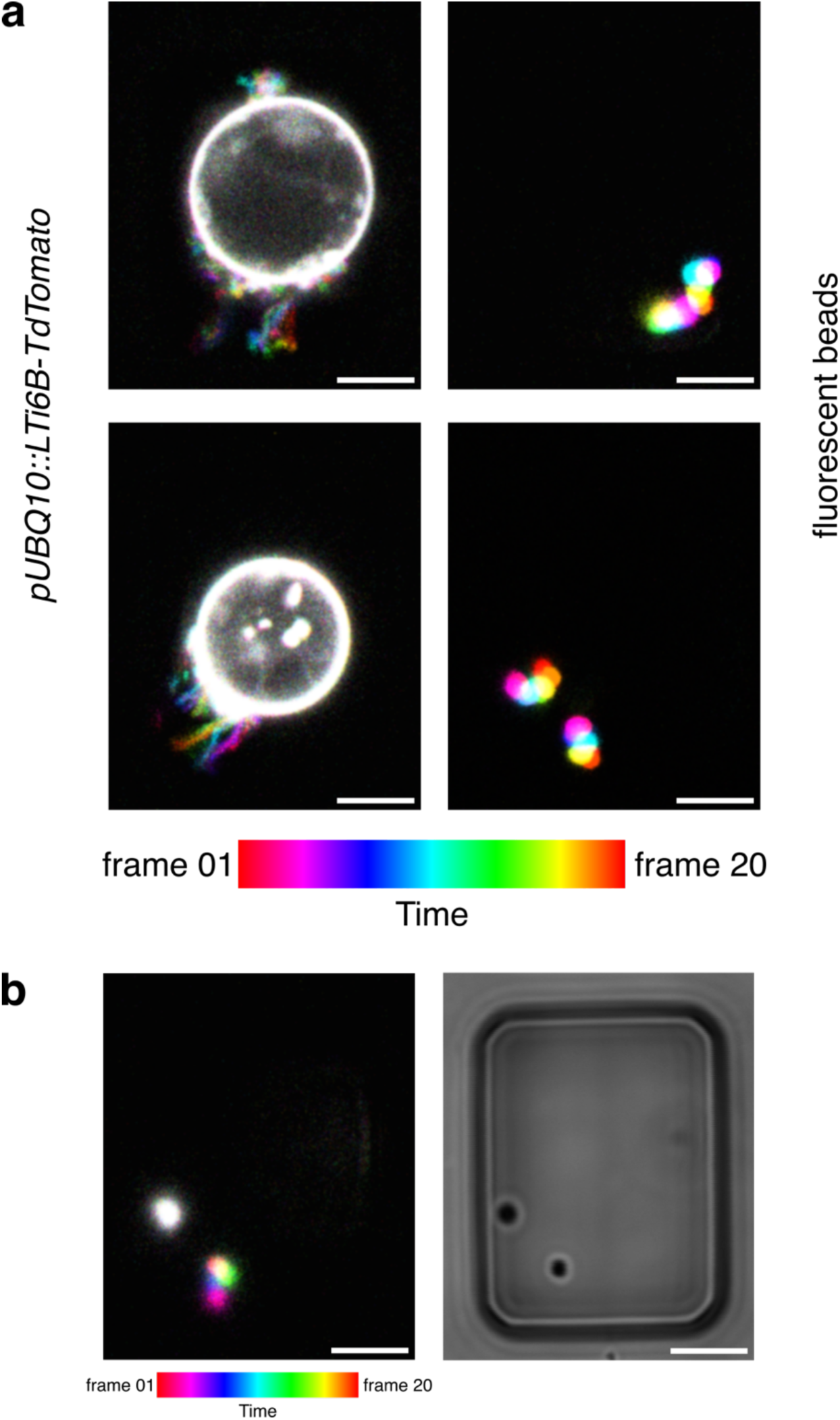
Additional examples of moving filopods and beads. **a**, The movement of the protrusions is qualitatively similar to the movement of fluorescent beads and hence may be passive. Timelapse of a single confocal plane of a protoplast with a plasma membrane marker (*pUBQ10::LTi6B-TdTomato*) (left) compared to fluorescent polystyrene beads (right), both in NOA73 microwells. 20 consecutive time frames (frame time 103ms) were color-coded and superposed to visualize movement, see also Movie S2. Images from the other two experimental repeats. **b**, Over time, the beads can get adsorbed to the walls of the microwells. Once adsorbed, they stop moving. Left, Bead movement represented as in **a**. Right, transmitted light image. Note that the bead adsorbed to the wall does not move (it appears white).

## Supplementary movies

**Movie S1: Filopods move dynamically.**

Timelapse movie of a protoplast in a NOA73 microwell with plasma membrane marker (*pUBQ10::LTi6B-TdTomato*) pseudo-colored in green, overlayed with chloroplast autofluorescence (pseudo-colored in magenta) and transmitted light. Fluorescent signal from single confocal plane. Frames are displayed at 7 fps and were acquired with a frame time of 630ms. Scale bar: 5 µm. The protoplast was obtained from 6-day old plants without cold stratification, using 1000 g for the centrifugation steps. A drop of 30 µl protoplast suspension was left to sink into the microwells for 1h and then imaged.

**Movie S2: The movement of the protrusions is qualitatively similar to the movement of fluorescent beads and hence may be passive.**

Dynamic representation of data displayed in Fig. 4b. Timelapse movie of a single confocal plane of a protoplast with a plasma membrane marker (*pUBQ10::LTi6B-TdTomato*) pseudo-colored in green (left) compared to fluorescent polystyrene beads (right) pseudo-colored in green, each overlayed with the transmitted light signal (note the microwell). Frames are displayed at 5 fps and were acquired with a frame time of 103ms.

**Movie S3-5: Filopods are occasionally observed to attach to the surface of the microwells.** Dynamic representation of data displayed in Fig. 5a. Examples from two different experiments: S3, S4: plasma membrane marker (*pUBQ10::LTi6B-TdTomato*), pseudo-colored in green; S5: tubulin marker (*p35S::GFP-TUA6*), pseudo-colored in green, each overlayed with transmitted light (note the microwell) and chloroplast autofluorescence (pseudo-colored in magenta). Fluorescence signal from single confocal plane. Scale bars: 5 µm. Frames are displayed at 7 fps and were acquired with a frame time of 103ms (S3,4) and 315ms (S5), respectively.

---

**Table S1:** Protoplast statistics (Fig. 1f): Summary of N = 3 experimental repeats with a total of n = 922 alive protoplasts.

| Experiment ID |  | smooth | filopods | dots |  |  |
| --- | --- | --- | --- | --- | --- | --- |
| PRO039 | Sum | 125 | 32 | 61 | Sum alive | 218 |
|  | Relative | 57% | 15% | 28% |  |  |
| PRO040 | Sum | 234 | 114 | 122 | Sum alive | 470 |
|  | Relative | 50% | 24% | 26% |  |  |
| PRO061 | Sum | 147 | 24 | 63 | Sum alive | 234 |
|  | Relative | 63% | 10% | 27% |  |  |
|  |  |  |  |  | Sum all alive | 922 |
| Mean relative |  | 57% | 16% | 27% | 1 |  |
| Standard deviation |  | 7% | 7% | 1% |  |  |

**Table S2:** Long-term survival rates of protoplasts with filopods, dots, and smooth protoplasts (Fig. 2b,c). Average and standard deviation of N = 3 independent experiments with 27, 35, 35 protoplasts, respectively.

| TOTAL |  |  | still alive after |  |  |
| --- | --- | --- | --- | --- | --- |
|  |  | 0h | 4h | 8h | 12h |
| filopods | average | 100 | 92% | 81% | 65% |
|  | stddev | 0 | 8% | 5% | 8% |
| dots | average | 100 | 94% | 83% | 83% |
|  | stdev | 0 | 10% | 29% | 29% |
| smooth | average | 100 | 90% | 74% | 66% |
|  | stdev | 0 | 18% | 24% | 30% |

**Table S3:** Fraction of smooth protoplasts and protoplasts with protrusions (dots + filopods) of “green” (*p35S::GFP-LTi6B*) and “magenta” (*pUBQ10::LTi6B-TdTomato*) protoplasts across N = 3 independent experiments with n = 324, 159, 91 alive protoplasts, respectively (Fig. 2d).

| Experiment ID | magenta smooth | green smooth | magenta + magenta protrusions | green + green protrusions | mixed protrusions |
| --- | --- | --- | --- | --- | --- |
| PRO043 | 62% | 67% | 38% | 33% | 0% |
| PRO045 | 65% | 72% | 35% | 28% | 0% |
| PRO059 | 66% | 56% | 34% | 44% | 0% |
| Average | 64% | 65% | 36% | 35% | 0% |
| Standard Deviation | 2% | 8% | 2% | 8% | 0% |

**Table S4:** Fractions of protoplasts with strong microtubule (MT) signal, some MT signal, only diffuse tubulin signal, sick/dead protoplasts (upper table, Extended Data Figure 8a). Fractions of protoplasts with strong MT signal in filopods, diffuse tubulin signal in filopods, diffuse tubulin signal in dots (lower table, Fig. 4a, Extended Data Figure 8b) for DMSO control and taxol treatment, respectively. N = 3 independent experiments with n = 98, 100, 100 protoplasts (DMSO) and n = 100, 100, 100 protoplasts (taxol).

**DMSO control**
|  | PRO055 | PRO066 | PRO082 | average | std dev |
| --- | --- | --- | --- | --- | --- |
| MTs | 5.10% | 6.00% | 14.00% | 8% | 5% |
| MT/diffuse | 45.92% | 39.00% | 53.00% | 46% | 7% |
| diffuse | 29.59% | 38.00% | 28.00% | 32% | 5% |
| sick/dead | 19.39% | 17.00% | 5.00% | 14% | 8% |
| MTs filopods | 1.02% | 0.00% | 2.00% | 1% | 1% |
| diffuse GFP-TUA6 signal in filopods | 5.10% | 3.00% | 6.00% | 5% | 2% |
| diffuse GFP-TUA6 signal in dots | 7.14% | 11.00% | 10.00% | 9% | 2% |
| SUM protrusions | 13.27% | 14.00% | 18.00% | 15% | 3% |
| SUM filopods | 6.12% | 3.00% | 8.00% | 6% | 3% |

**Taxol**
|  | PRO055 | PRO066 | PRO082 | average | std dev |
| --- | --- | --- | --- | --- | --- |
| MTs | 50.00% | 39.00% | 64.00% | 51% | 13% |
| MT/diffuse | 39.00% | 22.00% | 29.00% | 30% | 9% |
| diffuse | 3.00% | 13.00% | 1.00% | 6% | 6% |
| sick/dead | 8.00% | 26.00% | 6.00% | 13% | 11% |
| MTs filopods | 12.00% | 7.00% | 12.00% | 10% | 3% |
| diffuse GFP-TUA6<br>signal in filopods | 2.00% | 5.00% | 2.00% | 3% | 2% |
| diffuse GFP-TUA6<br>signal in dots | 5.00% | 3.00% | 4.00% | 4% | 1% |
| SUM protrusions | 19.00% | 15.00% | 18.00% | 17% | 2% |
| SUM(filopods) | 14.00% | 12.00% | 14.00% | 13.33% | 1.15% |

